# Jen1 transport and endocytosis in yeast reveal pH-responsive strategies to prevent metabolites loss

**DOI:** 10.1101/2024.08.01.606176

**Authors:** Cláudia Barata-Antunes, Marieke Warmerdam, Erik de Hulster, Inês P. Ribeiro, Clara Cardoso, Vítor Fernandes, Beatriz Leite, Margarida Casal, Hernâni-Gerós, Jack Pronk, Joaquín Ariño, Robert Mans, Sandra Paiva

**Affiliations:** Centre of Molecular and Environmental Biology, Department of Biology, University of Minho, 4710-057, Braga, Portugal; Institute of Science and Innovation for Bio-Sustainability (IB-S), University of Minho, 4710-057, Braga, Portugal; Department of Biotechnology, Delft University of Technology, Van der Maasweg 9, 2629 HZ, Delft, The Netherlands; Life and Health Sciences Research Institute (ICVS), University of Minho, Braga, Portugal; Centre for the Research and Technology of Agro-Environmental and Biological Sciences, University of Trás-os-Montes and Alto Douro, Vila Real, Portugal; Centre of Biological Engineering (CEB), Department of Biological Engineering, University of Minho, Braga, Portugal; Institut de Biotecnologia i Biomedicina & Departament de Bioquímica i Biologia Molecular, Universitat Autònoma de Barcelona, Cerdanyola del Vallès, Spain

**Author notes:** **Correspondence:** Sandra Paiva, Robert Mans.

**Keywords:** transporters, carboxylic acids, endocytosis, alkali stress, lactate, pH, chemostat cultivation

## Abstract

In batch cultures of *Saccharomyces cerevisiae* grown on lactate, medium alkalinization coincides with endocytosis of the Jen1 lactate/pyruvate transporter. To assess how culture pH impacts Jen1 endocytosis, *S. cerevisiae* was grown in carbon-limited continuous cultures with a mixed ethanol-lactate feed. When applying a linearly increasing pH (6.75 to 7.25), extracellular lactate and pyruvate concentrations increased progressively. Up to pH 7.0, these concentrations aligned with the thermodynamic equilibrium of Jen1-mediated electroneutral carboxylate-proton symport, while Jen1 internalization became more pronounced above pH 7.0. Transcriptional upregulation of genes involved in oxidative phosphorylation, together with higher residual lactate levels, indicated increased energy demands at supra-optimal pH values. By analyzing transport, endocytosis, and transcriptional responses in actively growing continuous cultures, this study uncovers pH-dependent physiological challenges associated with electroneutral carboxylate/proton symport. The data support the hypothesis that Jen1 internalization evolved to prevent intracellular metabolite loss under unfavorable pH conditions. These findings provide novel insights into the physiological consequences of transporter energy coupling and the ecophysiological relevance of anion transporters when extracellular pH approaches or exceeds cytosolic pH.

## 1. Introduction

In nature as well as in human-made environments, the yeast *Saccharomyces cerevisiae* faces changes in its surroundings. Cells can adapt to these changes, which include fluctuations in nutrient availability and extracellular pH, by reprogramming their metabolism, for instance by inducing or repressing transcription of specific genes [1–4]. Adaptation can also occur post-translationally, for example by promoting internalization (endocytosis) and degradation of plasma membrane (PM) transporters that are deleterious or dispensable under specific conditions [5–7].

Jen1 is a transmembrane transporter protein of the yeast *S. cerevisiae* that mediates symport of monocarboxylate anions such as lactate and pyruvate with a proton [8–11]. It is expressed and localized at the PM when *S. cerevisiae* is grown on non-fermentable carbon sources, including lactate, pyruvate, acetate and ethanol [12]. A variety of signals induce Jen1 internalization and degradation, including glucose [10, 13–19], rapamycin, and cycloheximide ([20] (see [5] for a review). When yeast cells are grown on monocarboxylates, they alkalinize their extracellular medium, which coincides with Jen1 endocytosis and degradation [20]. We showed that the Bul1 arrestin (an adaptor of Rsp5 ubiquitin ligase), an active TORC1 complex, and ammonium (a preferred nitrogen source for growth of *S. cerevisiae*), are necessary for efficient pH-dependent endocytosis of Jen1 [20]. However, the regulatory networks that connect alkalinization of the external medium to Jen1 downregulation remain elusive.

In shake-flask cultures, concentrations of biomass, substrate, oxygen as well as extracellular pH change during growth. All these changes can affect gene expression [21] and other cellular processes, including endocytosis of proteins localized at the PM. This complexity makes it difficult to unequivocally assess the impact of extracellular pH on Jen1-mediated carboxylate transport and on its endocytosis. In continuous bioreactor cultures, key culture parameters such as specific growth rate, temperature, biomass concentration, pH, and oxygen availability can be controlled independently and maintained at desired setpoints [22]. Use of continuous cultures therefore facilitates identification of the impact of individual culture parameters (e.g., extracellular pH) on gene expression [23].

The PM ATPase (Pma1) of *S. cerevisiae* cells generates an electrochemical proton gradient, the proton motive force (PMF), across the PM by pumping out a proton at the expense of ATP [24]. The PMF consists of the pH gradient (ΔpH = pH_in_ - pH_out_) and the electrical potential difference resulting from the charge gradient, generated by the movement of positive or negative charges to the extracellular space (Δ*ψ* = *ψ*_in_ – *ψ*_out_, in V) [25]. Because Jen1 mediates electroneutral monocarboxylate/proton symport, with a stoichiometry of 1 H^+^ (proton): 1 monocarboxylate (anion) [10], only the pH gradient (ΔpH) contributes to the driving force for accumulative monocarboxylate uptake [25]. In addition to its role in monocarboxylate uptake, Jen1 has been implicated in monocarboxylate export [26–29].

Facilitation of bidirectional transport implies that, when the external pH exceeds the internal pH, equilibrium of Jen1-mediated transport will occur at external substrate concentrations that are higher than those in the cytosol. We hypothesise that under these conditions, removal of Jen1 from the PM limits or prevents ‘leakage’ of Jen1 substrates from cells (**Fig. 1A**). To test this hypothesis and to further explore regulatory mechanisms underlying Jen1 endocytosis, *S. cerevisiae* was first grown in aerobic, carbon-limited continuous cultures on a mixture of ethanol and lactate. These continuous cultures were then subjected to a linear increase of the external pH, starting below and ending above pH 7.0, which based on literature is close to the cytosolic pH [30–33] of this yeast. Using this experimental set-up, we measured extracellular concentrations of the Jen1-transported metabolites lactate and pyruvate, as well as cellular localization of Jen1. Transcriptome profiles were then obtained from steady-state continuous cultures grown at selected pH values. Our results indicate that, with increasing pH, the reversible mode of action of Jen1 leads to increasing loss of pyruvate from the cells. At pH values above 7.0, Jen1 is largely internalized and this internalization is accompanied by transcriptional remodelling, seemingly aimed at improving respiratory capacity. These findings support the hypothesis that Jen1 internalization is a response to minimize the loss of essential metabolites when environmental pH surpasses intracellular pH.

**Fig. 1.**
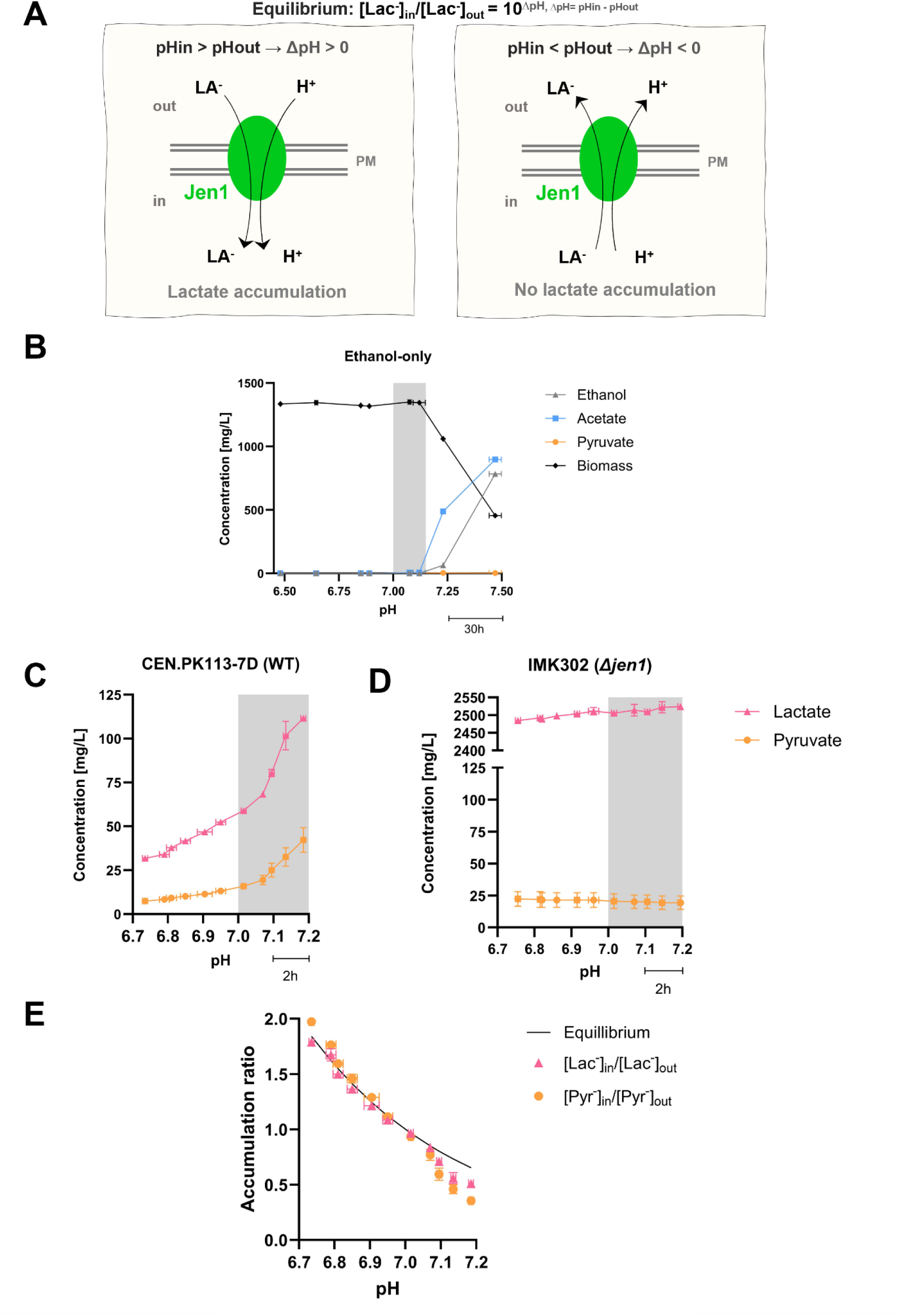
Physiology of *S. cerevisiae* strains in aerobic, carbon-limited continuous cultures subjected to a linear increase of extracellular pH. Continuous cultures grown at a dilution rate of 0.07 h^−1^ were continuously fed with SM containing 50 mM ethanol, either with or without 33 mM L-lactate. Cultivation experiments (panels B-E) were performed as independent duplicates and the data are presented as the mean, with error bars representing the standard error of the mean (SEM). **(A)** Schematic representation of the reversible ΔpH-dependent symport of lactate anions (La^−^) and protons (H^+^) by Jen1. At extracellular pH values below the cytosolic pH (ΔpH > 0), equilibrium of Jen1-mediated transport involves intracellular lactate accumulation, while the reverse situation occurs when ΔpH < 0. **(B)** Concentrations of biomass, ethanol and pyruvate in a continuous culture of *S. cerevisiae* CEN.PK113-7D grown on ethanol as sole carbon source and subjected to a linearly increasing pH (0.0083 ≈ 0.01 pH unit h^−1^). The time scale bar (30h) represents the duration required for each 0.25 pH unit increase. **(C)** Concentrations of lactate and pyruvate in a continuous culture of *S. cerevisiae* CEN.PK113-7D, fed with a mixture of ethanol and L-lactate and subjected to a linearly increasing pH (0.05 pH unit h^−1^). The time scale bar (2h) represents the duration required for each 0.1 pH unit increase **(D)** Concentrations of lactate and pyruvate in a continuous culture of a congenic *Δjen1 S. cerevisiae* strain, fed with a mixture of ethanol and lactic acid and subjected to a linearly increasing pH (0.05 pH unit h^−1^). **(E)** Accumulation ratios (AR, internal carboxylate concentration divided by external carboxylate concentration) of lactate and pyruvate during the pH ramp experiments shown in panel C and the corresponding independent replicate experiments. The solid black line indicates the AR calculated from the equilibrium equation (AR = 10^(pHin – pHout)^) and a pHin of 7.0. AR values were calculated from extracellular concentrations based on the assumption that intracellular concentrations of lactate and pyruvate concentrations were constant and independent of extracellular pH.

## 2. Results

### 2.1. Extracellular lactate and pyruvate concentrations in carbon-limited continuous cultures subjected to a linear pH ramp

A mixed-substrate continuous-cultivation system was implemented to study the impact of extracellular pH on activity and subcellular localization of the Jen1 monocarboxylate-proton symporter. Lactate was fed to these cultures along with ethanol, which is transported across the yeast PM by passive diffusion and neither represses *JEN1* promoter activity [12] nor interferes with Jen1 internalization and degradation.

To investigate up to which external pH value *S. cerevisiae* CEN.PK113-7D can grow on ethanol at a dilution rate of 0.07 h^−1^, continuous cultures grown on ethanol as sole carbon substrate were subjected to a pre-programmed pH ramp. When the culture pH was linearly increased from 6.50 to 7.50 at a rate of ≈0.01 pH units h^−1^, incomplete ethanol utilization occurred above pH 7.20 (**Fig. 1B** and **Table S1**). This observation and a progressive decrease of the biomass concentration in the cultures indicated that above pH 7.20, the specific growth rate declined to below 0.07 h^−1^.

Similarly, carbon-limited continuous cultures of *S. cerevisiae* CEN.PK113-7D fed with a mixture of 33 mmol L^−1^ L-lactate and 50 mmol L^−1^ ethanol (dilution rate 0.07 h^−1^) were first allowed to reach steady state at pH 6.75. After establishing steady-state growth, a pre-programmed pH ramp was initiated, during which the culture pH was linearly increased from pH 6.75 to 7.20 at a rate of 0.05 pH units h^−1^. During this pH ramp, extracellular concentrations of the Jen1 substrates lactate and pyruvate increased with increasing pH. This culture pH-dependent increase of extracellular lactate and pyruvate concentrations became more pronounced above pH 7.0 (**Fig.1C** and **Table S2**).

To investigate whether Jen1 was indeed responsible for the observed increase of extracellular pyruvate concentrations at increasing extracellular pH, additional pH-ramp experiment (pH 6.75 to pH 7.25, 0.01 pH unit h^−1^) was performed with a congenic *Δjen1* strain. In this experiment, no lactate consumption was observed, and extracellular pyruvate concentration did not increase with increasing pH (**Fig. 1D** and **Table S3**). These results are consistent with earlier reports showing that, in wild-type *S. cerevisiae*, Jen1 is the sole PM transporter for these monocarboxylates [9, 34].

When extracellular and intracellular pH are the same (ΔpH = 0), the equilibrium of Jen1-mediated electroneutral proton symport occurs at identical intra- and extracellular concentrations of its carboxylate substrates. Based on reported values of intracellular pH in *S. cerevisiae* [30–33, 35] we assumed this situation to occur at pH 7.0. Moreover, we hypothesise that, at the fixed specific growth rate in the continuous cultures, intracellular concentrations of lactate and pyruvate as well as intracellular pH were independent of culture pH. At extracellular pH values of 6.75 to 7.0, accumulation ratios of lactate and pyruvate, calculated based on these assumptions from measured extracellular concentrations, closely agreed with the theoretical equilibrium (internal/external concentration = 10^ΔpH^ when pK_a_ << pH_out_, pH_in_; **Fig. 1E**). As anticipated, calculated accumulation ratios for lactate and pyruvate became smaller than 1 at pH values above 7.0. However, at those pH values, accumulation ratios were even lower than predicted based on the abovementioned assumptions. This observation may reflect that, at external pH values above 7.0, elevated intracellular concentrations of lactate and pyruvate are needed for sustaining higher rates of lactate dissimilation in order to generate additional ATP for countering alkaline stress.

### 2.2 Localization and internalization of Jen1 in response to a linear increase of extracellular pH

In contrast to lactate, pyruvate was not included in the feed of the continuous cultures. The observed extracellular pyruvate concentrations in the pH-ramp experiments therefore reflected net Jen1-mediated export of pyruvate from the cells. Preventing release of pyruvate, a central intermediate in yeast metabolism [36], might provide a rationale for the previously reported removal of Jen1 from the PM upon exposure to alkaline pH in shake-flask cultures [20]. To specifically investigate the influence of extracellular pH on subcellular localization of Jen1, a pH-ramp experiment (pH 6.5 to 7.5, 0.01 pH unit h^−1^) was performed with continuous cultures of a congenic *S. cerevisiae* strain CB270 that constitutively expresses a Jen1-GFP fusion protein (**Fig. 2**).

**Fig. 2.**
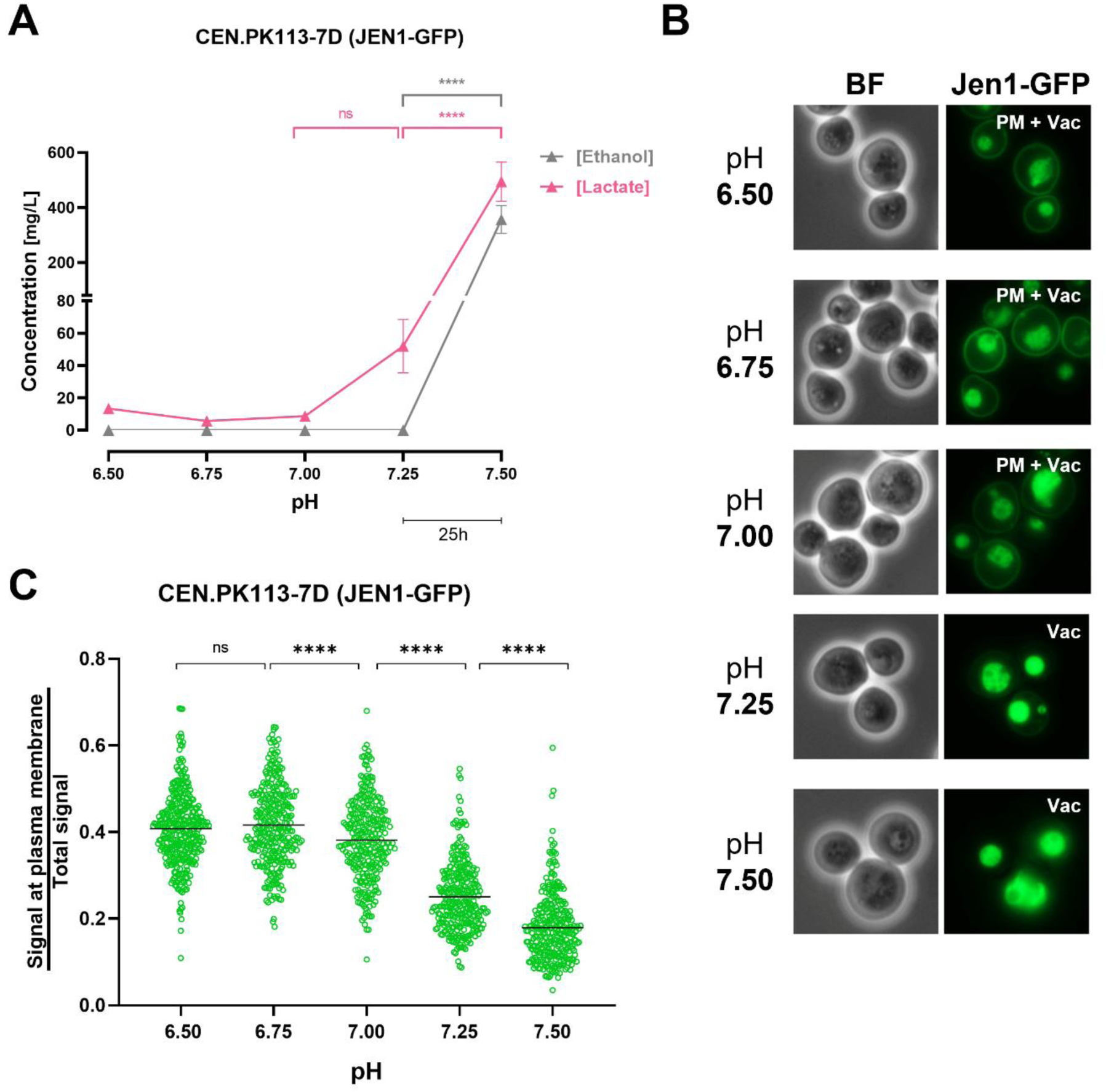
Evaluation of Jen1-GFP localization in *S. cerevisiae* CB270 (*JEN1::GFP::HghMX*) grown in aerobic, carbon-limited continuous cultures subjected to a linear increase of extracellular pH. Duplicate continuous cultures grown at a dilution rate of 0.07 h^−1^ were continuously fed with SM containing 50 mM ethanol and 30 mM L-lactate and subjected to a linearly increasing pH ramp (0.01 pH units h^−1^). **(A)** Extracellular concentrations of ethanol and lactate (mg L^−1^). Data are presented as the means of two individual experiments, with error bars representing the standard error of the mean (SEM). ns, not significant (*P*=0.8095); \*\*\*\**P*<0.0001 (two-way ANOVA analysis with a Tukey’s multiple comparisons test). The time scale bar (25h) represents the duration required for each 0.25 pH unit increase **(B)** Visualization of Jen1-GFP by epifluorescence microscopy at the indicated extracellular culture pH. The primary localization of the Jen1-GFP is described at the top left corner of each fluorescence microscopy image. PM, plasma membrane; Vac, vacuole; BF, bright field. **(C)** Quantitative analysis of the ratio of peripheral fluorescence over total fluorescence. Data are represented as scatter plot (n ≥ 300 cells for each plot), median values are indicated by horizontal lines. ns, not significant (*P*>0.05); \*\*\*\**P*<0.0001 (ordinary one-way ANOVA analysis with Tukey’s multiple comparisons test).

Culture samples were collected at every 0.25 pH interval for lactate and acetate measurements, as well as microscopy analysis. The residual concentration of lactate in continuous cultures did not significantly change between pH 6.5 and 7.0 (**Fig. 2A** and **Table S4**). However, above pH 7.0, the lactate concentration steeply increased from 0.009 ± 0.001 g/L (mean ± s.d.) to 0.495 ± 0.101 g/L at pH 7.5. No extracellular ethanol was detected when cultures were grown at pH values between 6.5 and 7.25, but at pH 7.5 an extracellular concentration of 0.36 ± 0.07 g L^−1^ ethanol was measured, coincident with a decrease in biomass dry weight (**Table S5**). This observation is consistent with results from cultures grown on ethanol as sole carbon source (**Fig. 1B**), in which cells were unable to keep up with the dilution rate of the continuous cultures (0.07 h^−1^) at pH values above 7.25.

Fluorescence microscopy (**Fig. 2B**) clearly revealed presence of Jen1-GFP at the PM at culture pH values of 6.5, 6.75 and 7.0. In contrast, at pH 7.25 and 7.50, visual inspection no longer revealed fluorescence at the PM and, instead, showed a predominant localization of fluorescence in the vacuole (**Fig. 2B**). In line with these visual observations, the ratio of peripheral fluorescence over total fluorescence by image quantitative analysis showed a decrease from 0.42 ± 0.091 (mean ± s.d.) at pH 6.75 to 0.18 ± 0.077 at pH 7.5 (**Fig. 2C**). These results demonstrate that the threshold extracellular pH at which Jen1-GFP is removed from the PM lies between pH 7.0 and 7.2.

### 2.3 Extracellular metabolite concentrations and Jen1 internalization in steady-state cultures grown at different extracellular pH values

Due to their dynamic nature, the pH-ramp experiments do not necessarily reflect the physiology of cells that are fully adapted to a specific extracellular pH. Therefore, based on results from the pH-ramp experiments, duplicate steady-state continuous cultures (dilution rate, 0.07 h^−1^) of the Jen1-GFP expressing strain CB270 were grown on ethanol/lactate at four different culture pH values: pH 6.0, pH 6.5 (Jen1 at PM in pH ramp experiments), pH 7.0 and pH 7.1 (close to Jen1 internalization threshold pH identified in pH-ramp experiments). Upon reaching steady state, culture samples were taken for metabolite measurements, microscopy and transcriptome analysis.

Attempts to establish stable steady-state cultures at pH 7.2, where Jen1 was largely removed from the PM in the pH-ramp experiments, were unsuccessful due to culture wash-out. This indicated that cells were unable to sustain a specific growth rate that matched the dilution rate (0.07 h^−1^). This result was consistent with results from pH-ramp experiments with ethanol-grown cultures of strain CEN.PK113-7D (**Fig. 1B**), in which pH 7.2 coincided with the point at which extracellular ethanol and acetate started to appear.

Over the evaluated range of culture pH values, no extracellular ethanol was detected, indicating that the steady-state cultures were able to maintain a specific growth rate of 0.07 h^−1^. The residual concentration of lactate in the cultures, which was approximately 13 mg L^−1^ at pH 6.0 and 6.5, and 30 mg L^−1^ at pH 7.0, increased to 92 mg L^−1^ at pH 7.1 (**Fig. 3A**; see also **Table S6**).

**Fig. 3.**
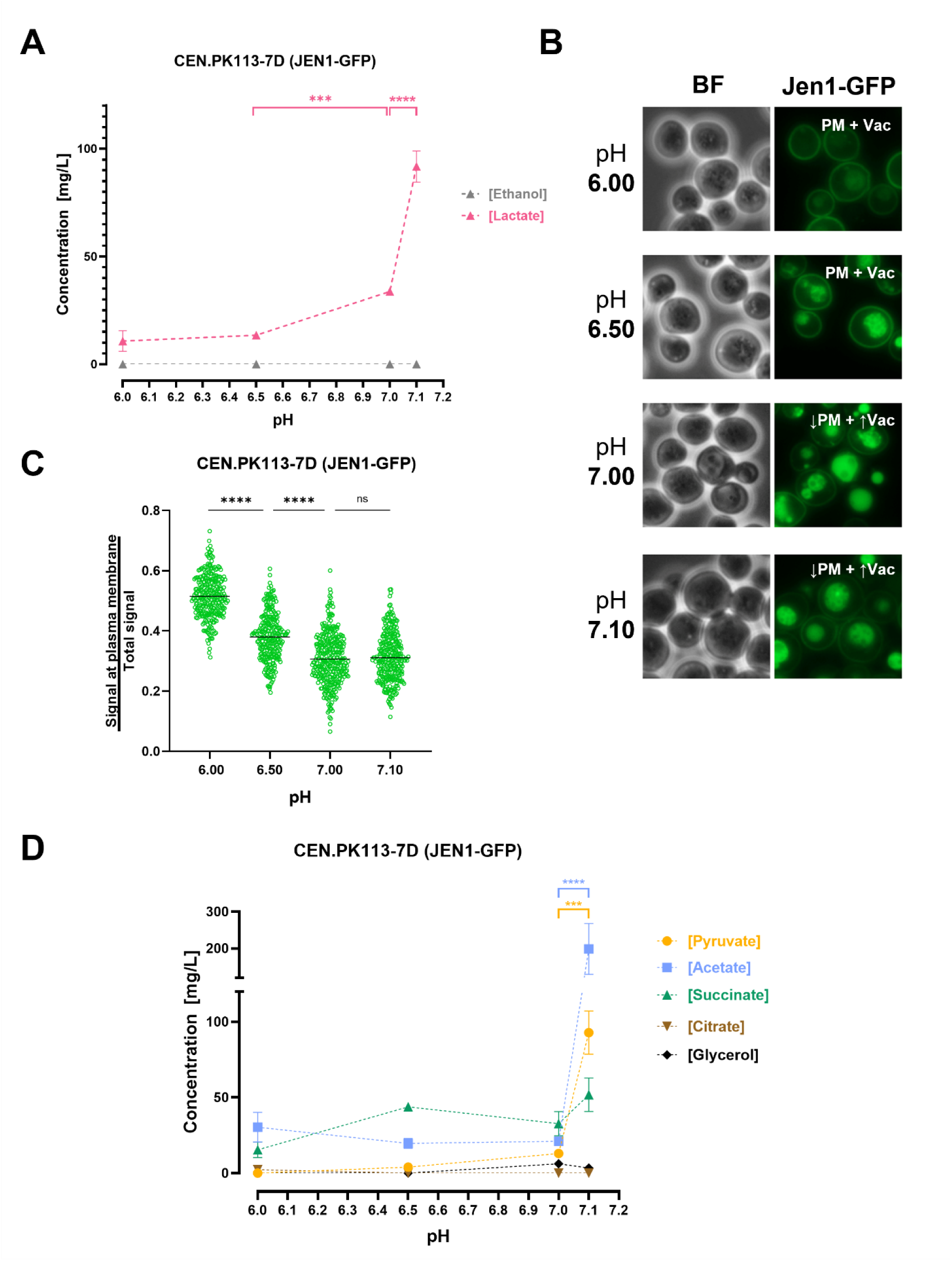
Extracellular metabolite concentrations and Jen1 internalization in steady-state continuous cultures grown at different extracellular pH values. Steady-state, carbon-limited continuous cultures of *S. cerevisiae* CB270 (CEN.PK113-7D *JEN1::GFP::HghMX*) grown at a dilution rate of 0.07 h^−1^ were fed with SM containing 50 mM ethanol and 30 mM L-lactate. Culture pH was maintained at 6.0, 6.5, 7.0, 7.1 or 7.2 by automated addition of 2 M KOH or 2 M H2SO4 (see Material and Methods). **(A)** Extracellular concentrations of ethanol and lactate (mg L^−1^). Data are presented as the means of at least two individual experiments, with error bars representing the standard error of the mean (SEM), \*\*\**P*=0.0004; \*\*\*\**P*<0.0001 (two-way ANOVA analysis with Tukey’s multiple comparisons test). **(B)** Visualisation of Jen1-GFP by epifluorescence microscopy at the indicated extracellular culture pH. The primary localization of Jen1-GFP is at the top left corner of each fluorescence microscopy image. PM, plasma membrane; Vac, vacuole; BF, bright field. **(C)** Quantitative analysis of the ratio of peripheral fluorescence over total fluorescence. Data are represented as scatter plot (n ≥ 270 cells for each plot), median values are indicated by horizontal lines. ns, not significant (P>0.05); ****P<0.0001 (ordinary one-way ANOVA analysis with Tukey’s multiple comparisons test. **(D)** Extracellular concentrations of metabolites other than lactate and ethanol (mg L^−1^). Data are presented as the means of at least two individual experiments, with error bars representing the standard error of the mean (SEM), \*\*\**P*=0.0006; \*\*\*\**P*<0.0001 (two-way ANOVA analysis was performed, followed by a Tukey’s multiple comparisons test).

Fluorescence microscopy of cells from steady-state cultures grown at pH 6.0 and 6.5 showed localization of Jen1-GFP at the PM and in the vacuole (**Fig. 3B**). In cultures grown at extracellular pH values of 7.0 and 7.1, less Jen1-GFP was found at the PM, while the fluorescence in the vacuole increased. In line with these observations, image analysis yielded a peripheral/total fluorescence ratio of 0.51 ± 0.07 at an extracellular pH of 6.0 and of approximately 0.3 at extracellular pH values of 7.0 and 7.1 (**Fig. 3C**).

In addition to the extracellular concentrations of lactate, also those of pyruvate, acetate, succinate, citrate, and glycerol were determined in culture supernatants of each steady-state culture (**Fig. 3D**, **Table S7**). These analyses indicated a 7- to 9-fold difference in the extracellular concentrations of pyruvate and acetate between cultures grown at pH 7.0 and pH 7.1. No such pronounced differences were observed for the other metabolites.

### 2.3 Transcriptome sequencing (RNA-seq) analysis of steady state-, aerobic-, continuous cultures of *S. cerevisiae* grown at different extracellular pH values

After defining the pH range at which Jen1 localized to the vacuoles, we aimed to investigate the metabolic status of the cell at different pHs and identity the pathways potentially involved in Jen1 internalization. To achieve this, samples of steady-state, aerobic continuous cultures of *S. cerevisiae* grown at pH values of pH 6.0, 6.5, 7.0, and 7.1 were collected for RNA sequencing. The RNA-seq analysis data was combined, and comparisons were made between the datasets obtained at the different pH values. Because the cells being compared were exposed to relatively small pH changes, we set a relatively low-stringency threshold (log_2_ >0.6 or <-0.6, with p-value < 0.01) to select for differentially expressed genes (DEGs). As expected, the effect on the number of DEGs was related to the intensity of the pH stress. Thus, the increase of pH from 6.0 to 6.5 affected a relatively small number of genes, barely above 100 (most of them induced, **Fig. 4** and **Fig. S1**). A further increase of the culture pH to 7.0 resulted in a higher number of genes with altered expression levels, with a particular increase in the number of those whose mRNA decreased. Interestingly, a relatively small increase of the culture pH by 0.1 units (7.1vs6.0 in comparison with 7.0vs6.0) doubled the number of mRNAs that were altered, with a nearly 3-fold increase in the number of repressed genes. Comparison of the data obtained at pH 7.0 with that found at pH 7.1 (**Fig. 4A**) yielded 108 genes with altered expression (mostly repressed). Considering that this change in pH results in a relatively minor alteration in proton concentration in comparison with the other changes examined, this transcriptional response can be considered meaningful.

**Fig. 4.**
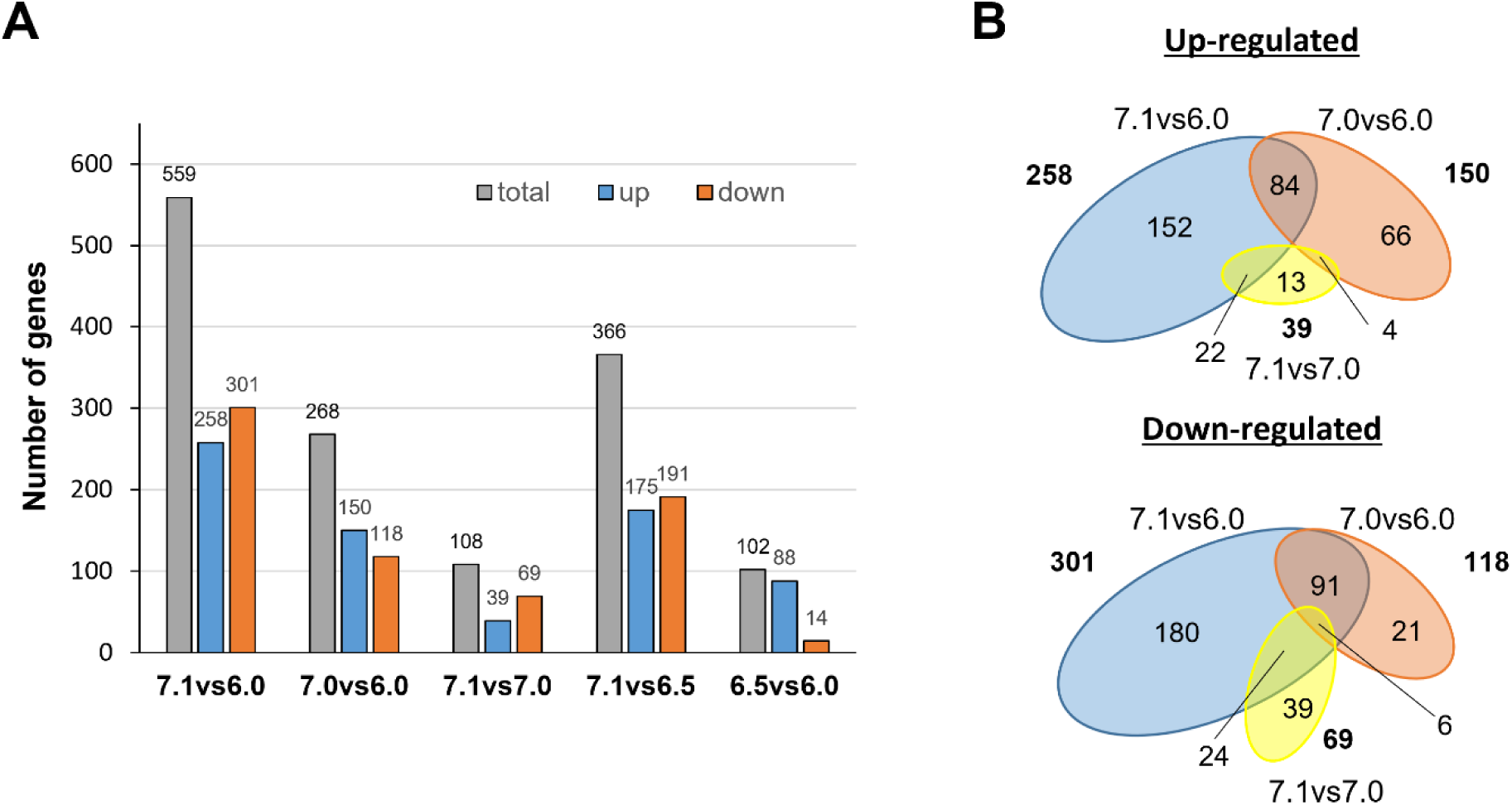
Transcriptional impact of external pH in steady-state continuous cultures of the *S. cerevisiae* strain CB270. Continuous cultures were grown at a dilution rate of 0.07 h^−1^ and fed with SM containing 50 mM ethanol and 30 mM L-lactic acid at pH 6.0, pH 6.5, pH 7.0 and pH 7.1. **(A)** Pairwise comparison of transcriptome data from cultures grown at different pH values (6.5vs6.0; 7.0vs6.0; 7.1vs6.0; 7.1vs6.5 and 7.1vs7.0). The grey column represents the total number of altered genes, and the blue and orange columns represent the number of up-regulated or down-regulated genes, respectively. **(B)** Venn diagrams of differentially expressed genes (DEG) for the pH 7.1vs6.0 (blue), 7.0vs6.0 (orange), and 7.1vs7.0 (yellow) data sets. The number of genes in each dataset are denoted in bold and is proportional to the area of the ellipses. The overlapping region indicates the numbers of DEGs that are common in the compared groups.

The transition from pH 6.0 to pH 7.0 or 7.1 resulted in changes consistent with previously reported data from cells subjected to mild alkalinization (see Discussion). Thus, diverse genes related to iron uptake and metabolism (*SIT1*, *FIT2*, *FIT3*, *FRE1*, *FET3*, and *HXM1*), high-affinity phosphate (*PHO84*, *PHO89*), or hexose transport (*HXT14*, *HXT10*, *HXT2*) were induced by the shift to either pH 7.0 or 7.1 (**Fig. S2**). **Figure 4B** shows the overlap of up-regulated or down-regulated genes between the following pH comparisons 7.1vs6.0, 7.0vs6.0 and 7.0vs7.1. It is worth noting that 174 of the 258 genes (59.7%) induced when comparing pH 7.1 and 6.0 were not found to be induced when comparing pH 7.0 and 6.0, whereas 204 out of 301 (67.8%) were found repressed in the former case but not in the latter. This suggests that the transition from pH 7.0 to pH 7.1, although it might appear quantitatively minor, has a notable impact in the expression profile of the cells. Since the most pronounced internalization of Jen1 seemed to be triggered by events during this transition, we found it pertinent to further examine the 174 genes specifically induced in cells exposed to the pH change from 6.0 to 7.1, but not from 6.0 to 7.0. As shown in **Fig. 5A** and **Fig. S3A**, this set of 174 induced genes was strongly enriched for genes related to mitochondrially-related events, with 70 genes (40%) encoding mitochondrial membrane-associated proteins (**Table S8**). Thirty-three genes were directly related to mitochondrial translation, including those encoding ribosomal proteins (such as *MRPL17*, *MRPL3*, *MRPL10*, and others), translation factors (such as *MRF1*, *MEF1*, *IFM1*, or *TUF1*), or RNA aminoacyl ligases (including *SLM5*, *MSR1*, *MST1*, *ISM1* and others). It is worth noting that genes encoding cytosolic ribosomal genes were not induced at all (**Fig. 5A**, **Fig. S3A**). In addition, sixteen genes were related to oxidative phosphorylation, either as members of the electron transport chain (such as *RIP1*, *SDH3*, *COX6*, COX*10*, *COX11* and others) or as components of the ATP synthase complex (*ATP2*, *ATP4*, *ATP5*, *ATP16*, *ATP17*, and *TIM11*). Additionally, substantial downregulation of mitochondrially-related genes is observed in the shift from pH 6.0 to 7.0, but not when pH changes from 6.0 to 7.1 (**Fig. S3B**).

**Fig. 5.**
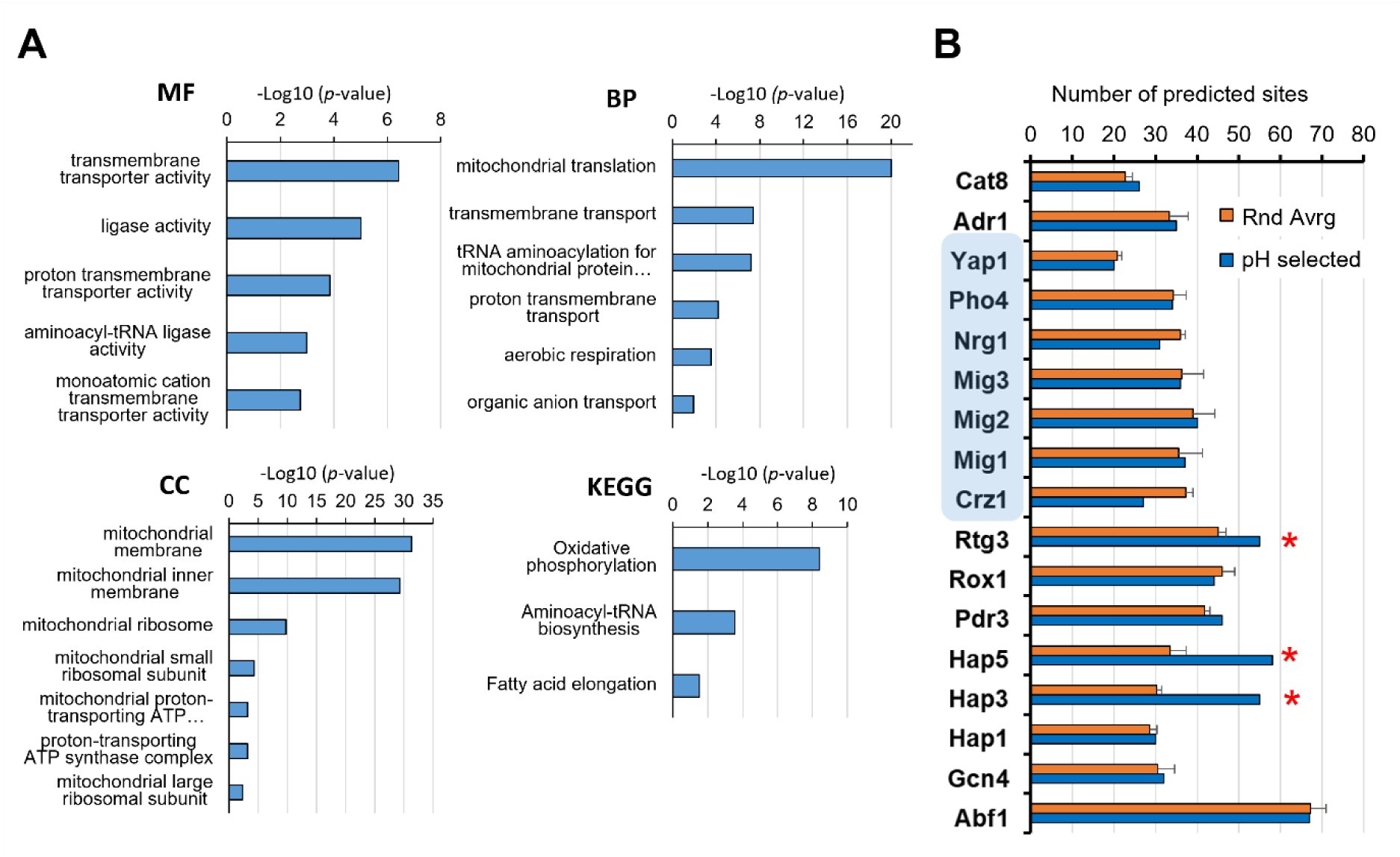
Gene Ontology and promoter analysis of the 174 genes specifically induced at the pH 6.0 to 7.1 transition. **(A)** The GO terms of the MF (Molecular Function), BP (Biological Process), and CC (Cellular component) domains enrichment of the 174 differentially expressed (up-regulated) genes. GO, Gene Ontology; KEGG, Kyoto Encyclopaedia of Genes and Genomes biological pathways database. The *p*-value threshold was set at 0.05. **(B)** The JASPAR database matrices were used to scan for transcription factor binding sites in the promoters of the mentioned genes (pH selected). As reference, four sets of 174 randomly selected genes were also scanned and the mean ± SEM of the number of predicted sites is offered for comparison. The asterisks denote binding sites whose abundance is markedly increased in the pH selected gene cohort. The blue shading denotes transcriptions factors whose activity is affected by alkalinization according to the literature (see main text).

The promoter region of the 174 genes was investigated for predicted transcription factor binding sites. As shown in **Fig. 5B**, such promoter regions were not enriched for consensus sequences corresponding to activators or repressors (shaded in blue) that, according to the literature [37–39], are known to play a relevant role in the overall adaptation to alkalinization. In fact, a decrease for the calcium-calcineurin activated Crz1 transcription factor was observed. In contrast, an enrichment for Rtg3 and, in particular, for Hap3 and Hap5 was detected. Rtg3 is a positive regulator of the retrograde pathway, which responds to mitochondrial dysfunction by adapting cell metabolism to the loss of tricarboxylic acid (TCA) cycle activity ([40, 41], reviewed in [42]). Hap3 and Hap5 are subunits of the Hap2p/3p/4p/5p CCAAT-binding complex, a transcriptional activator of respiratory gene expression (TCA cycle and oxidative phosphorylation genes) [43, 44]. Interestingly, the minimal change from pH 7.0 to 7.1 resulted in the transcriptional upregulation of 39 genes, of which only four (the permeases *DIP5*, *BAP3*, *GNP1*, and *TAT1*) were also induced in the transition from pH 6.0 to 7.0. Gene Ontology analysis of the 35 non-induced genes yielded a clear enrichment (*p*-value=1.8E-6) in genes encoding components of the mitochondrial inner membrane thus reinforcing the notion that the transition from pH 7.0 to 7.1 has a considerable impact on the regulation of mitochondrial function.

## 3. Discussion

Electroneutral symport of a proton and a monovalent carboxylate anion across the yeast plasma membrane, as mediated by Jen1, allows for ΔpH-driven accumulative anion import when the external pH is lower than the intracellular pH. For example, at an external pH of 5, which is near the optimum pH for growth of *S. cerevisiae* [45] and at a cytosolic pH of 7, equilibrium of Jen1-mediated lactate transport occurs at an intracellular-to-extracellular accumulation ratio of 100 of the anion. Correcting for the pK_a_ of lactic acid of 3.86 (see [46] for equation), this corresponds to a combined lactate/lactic acid accumulation ratio of 93-fold. However, as the extracellular pH increases, the contribution of ΔpH to the driving force for lactate uptake decreases. When the external pH becomes higher than the cytosolic pH, this contribution even becomes negative. The results presented in this study provide new insights into the physiological consequences of the mode of energy coupling of Jen1-mediated monocarboxylate transport and into the ecophysiological relevance of Jen1 endocytosis in situations where the extracellular pH equals or exceeds the cytosolic pH.

At a constant cytosolic pH, the equilibrium of Jen1-mediated transport ([lactate]_in_/[lactate]_out_ = 10^ΔpH^) dictates that, as the external pH increases, maintenance of a constant intracellular lactate concentration requires the extracellular lactate concentration to increase. Such an increase of the extracellular lactate concentration was indeed observed in continuous cultures grown on a lactate-ethanol mixture that were subjected to a linearly increasing extracellular pH (**Fig. 1**). Moreover, extracellular lactate concentrations measured in those cultures were consistent with Jen1-mediated transport operating at or close to thermodynamic equilibrium. This implies that, under the experimental conditions, *in vivo* activity of Jen1 did not constrain lactate transport across the PM. In natural environments with near-neutral or mildly alkaline conditions, the associated higher extracellular lactate concentrations are likely to negatively affect competition for lactate with microorganisms that employ Δψ- or ATP-coupled transporters and, consequently, can achieve much higher accumulation ratios. Based on our experiments with the Jen1 model system, we therefore hypothesize that electroneutral anion-proton symport is likely to predominantly occur in microorganisms adapted to environments with a pH below 7.

In the pH-ramp experiments shown in **Fig. 1**, calculated accumulation ratios of pyruvate, for which Jen1 is the only known efficient transporter in wild-type *S. cerevisiae* [34], at different external pH values, were virtually identical to those of lactate (**Fig. 1E**). Equilibration of in- and extracellular concentrations of essential intracellular metabolites such as pyruvate can potentially have a large impact on microbial cells, as illustrated by a simple example. One gram of yeast biomass has an intracellular volume of approximately 2 mL [47]. In a 1-L culture containing 100 mg yeast biomass L^−1^, intracellular volume accounts for just 0.2% of the total culture volume. Assuming an intracellular pyruvate concentration of 1 mmol L^−1^, equilibrium of Jen1-mediated transport at ΔpH = 0 will result in a situation where the biomass contains 0.2 μmol pyruvate, while the external medium contains 1 mmol pyruvate (with a mass of 88 mg, equal to 88% of the total cell mass in the culture). In other words, the cells will export their total weight in the form of pyruvate before equilibrium of transport over the membrane is reached. As illustrated in **Fig. 1** and **Fig. 3**, the extent of this metabolite loss will become even more pronounced at pH values above 7. The metabolic burden of symporter-mediated electroneutral carboxylate efflux can be even further enhanced when, in natural environments, biomass concentrations are lower, or when competing microorganisms and/or dilution continuously decrease extracellular concentrations of the ‘leaked’ carboxylate. Although *S. cerevisiae* is clearly not optimally adapted for growth at neutral or mildly alkaline environments, this does not rule out an evolutionary significance of preventing metabolite leakage during transient exposure to such conditions. Use of continuous cultures, in which specific growth rate and other growth parameters were carefully controlled, enabled unequivocal identification of extracellular pH as a key trigger for removal of Jen1 from the plasma membrane. The maximum culture pH of approximately 7.2, imposed by the continuous cultivation set-up, did not allow us to capture the complete internalization of Jen1 that was previously observed in batch cultures [20], which demonstrated complete Jen1 internalization. However, in cultures grown at pH 7.0 and 7.1, the majority of a Jen1-GPF fusion protein was found in the vacuole. This result supports the hypothesis that Jen1 internalization has evolved as a mechanism to prevent intracellular metabolite loss during exposure to alkaline environments.

Transcriptome data corresponding to the transitions from pH 6.0 to pH 7.0 or 7.1 reveals an increase in mRNA levels of genes involved in iron and copper uptake and metabolism, as well as in high-affinity phosphate and sugar transport. These changes fit well with previously reported data that different research groups obtained from exponentially growing batch cultures that were subjected to moderate alkalinization [37, 48, 49]. However, our data also show that the transition to pH 7.1 triggers a specific response that is not observed in cells shifted from pH 6.0 to 7.0. This response involves induction of genes involved in oxidative phosphorylation (**Fig. 6**) and suggests that growth at pH values above 7.0 requires increased rates of respiration-coupled ATP generation. At those pH values, extracellular concentrations of lactate and pyruvate in pH-ramp experiments could no longer be fitted based on an assumed strict homeostasis of intracellular lactate and pyruvate concentrations (**Fig. 1**). In line with the transcriptome data, the higher inferred intracellular lactate and pyruvate concentrations in cultures grown at pH values above 7.0 (**Fig. 1E**) may reflect increased lactate dissimilation enabled by elevated intracellular substrate concentrations of key enzymes and metabolites in the dissimilatory pathway. Our previous study demonstrated that Jen1 internalization, triggered by the prolonged growth on lactate and consequent alkalinization, seemed to involve the Bul1 α-arrestin and the TORC1 pathway [20]. However, the transcriptomic data obtained from continuous cultures in this study did not reveal an upregulation of the TORC1 pathway when pH 6.0 to 7.1 conditions were compared (**Fig. 5**). The differences in growth conditions between these studies, where the continuous cultures in this work were exposed for longer periods to the alkaline conditions, could have led to the use of an alternative intracellular pathway for Jen1 degradation.

**Fig. 6.**
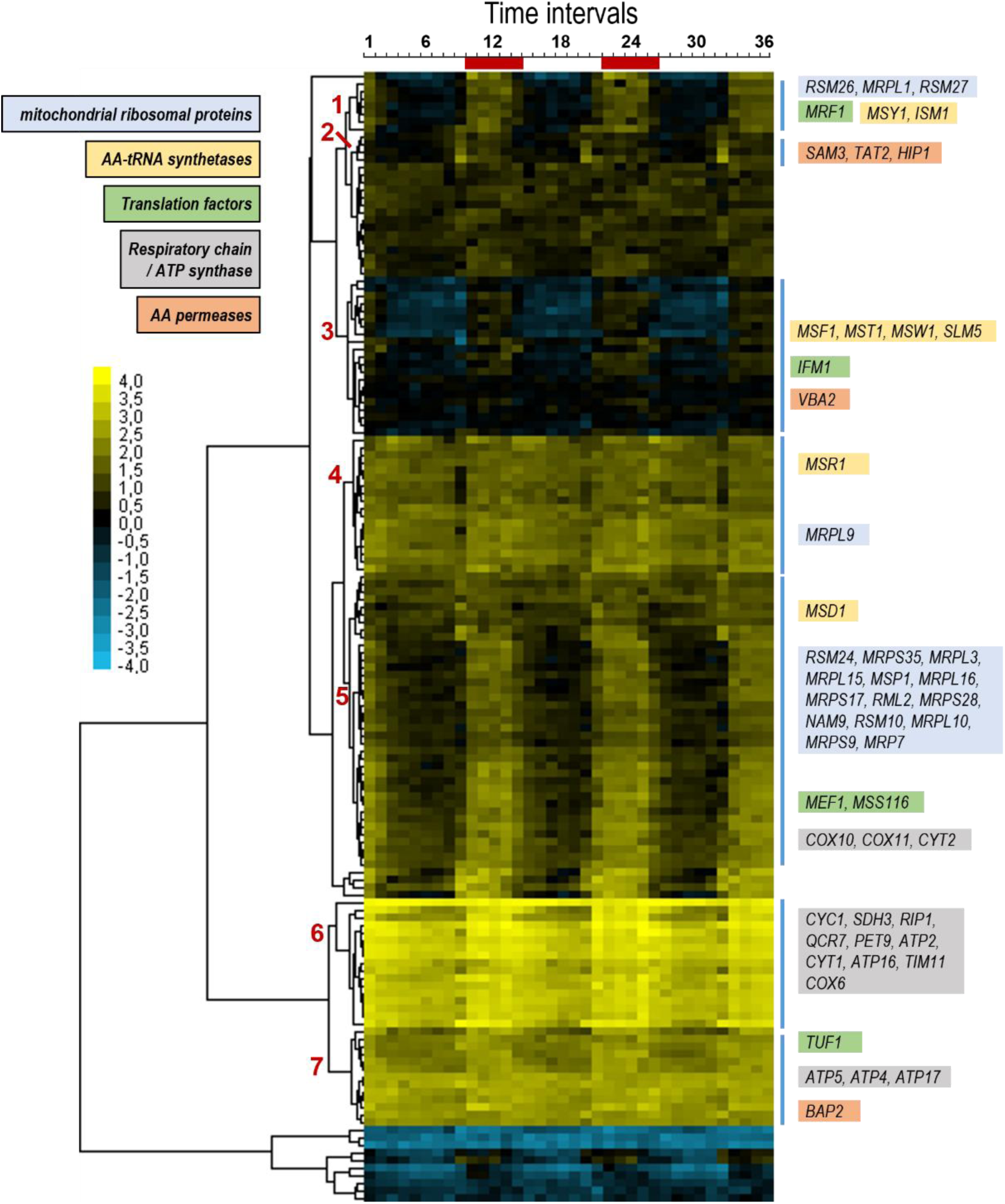
Periodic profile of genes specifically induced at the pH 6.0 to 7.1 transition. Our list of 174 genes was crossed with the expression data provided by Tu and coworkers [50] which described the cyclic expression profile of yeast cells grown under continuous, nutrient limiting conditions. The data for the 149 genes common to both datasets was clustered with the Cluster 3.0 software (Euclidean distance, average linkage) [56] and plotted with JavaTreeView [57]. The numbers on the left identify the main clusters, with relevant genes shown on the right according the indicated functional colour code. Time intervals at the top correspond to the interval of sampling (approximately 25 min). The red bars show the time intervals corresponding to the reductive/building (R/B) period (see main text and reference [50]).

Tu *et al* [50] showed that *S. cerevisiae* cells grown under continuous, nutrient-limited conditions, after a short initial starvation period, exhibit highly periodic cycles related to respiratory bursts. They identified three repetitive superclusters, oxidative (Ox), reductive/building (R/B), and reductive/charging (R/C), thus defining three major phases of the yeast metabolic cycle. We soon realized that the top 40 R/B genes identified, showing stronger periodicity, included 13 of our 174 genes, whereas none of them appeared in the top 40 Ox or R/C lists. To better quantify this apparently specific correspondence, we performed a cluster analysis with the transcriptomic data reported in [50] for the 149 genes in common with our 174 gene list. As presented in **Figure 6**, with few exceptions all these genes are expressed following the R/B supercluster pattern. As the mentioned R/B supercluster peaks when cells are about to cease oxygen consumption, it could be interpreted as a preparatory step to meet an acute requirement to increase respiratory capacity. It must be noted that the expression of many of the genes identified in the specific switch from pH 6.0 to 7.1 does not seem linked to a general response to alkalinization, as the promoters of these genes are not enriched in binding sites for any of the diverse transcription factors known to shape the alkaline pH response [39]. Instead, such response is clearly connected to the mitochondria and appear to indicate a first step in intensifying the capacity for mitochondrial respiration. This is reflected in the enrichment in Rtg3 and, specially, Hap3 and Hap5 binding sites in their promoters (**Fig. 5B**), as these transcription factors are pivotal components of the adaptation to respiration [44]. Interestingly, an increase in the TCA cycle activity and respiratory rate when batch cultures of yeast cells were switched to pH 7.5 was reported [51].

Metabolite analyses, subcellular localization studies and transcriptome analysis on carefully controlled continuous cultures subjected to defined pH changes enabled a deeper understanding of the mode of action, regulation and disadvantages of carboxylate transport by Jen1 at neutral and alkaline pH. It will be interesting to apply the same approach to investigate pH-dependent activity and localization of additional H^+^/carboxylate symporters in *S. cerevisiae* and other eukaryotes. Interesting targets for such studies include the *S. cerevisiae* H^+^-uracil symporter Fur4, which has been implicated in loss of intracellular uracil under alkaline conditions [52] and acetate transporters, as acetate is released by ethanol-grown (**Fig. 1B**) and glucose-grown [53] *S. cerevisiae* cultures at extracellular pH values above 7 and the scientific literature proposes different energy-coupling mechanisms (reviewed in [54, 55]).

## 4. Material and Methods

### 4.1. Yeast strains and storage conditions

The *S. cerevisiae* strains used in this study are listed in **Table 1**. For prolonged storage, the strains were grown at 30 °C, 200 rpm, in YPD (1 % (w/v) yeast extract, 1 % (w/v) peptone and 2 % (w/v) glucose) medium. After overnight growth, glycerol was added to a final concentration of 30 % (v/v) and 1 mL aliquots were stored at – 80 °C.

**Table 1.** List of the strains used in this work.

| Strain | Genotype | Reference/source |
| --- | --- | --- |
| Ace55 | W303-1A *JEN1::GFP::KanMX leu2 ura3 trp1 his3 ade2* | Our laboratory collection |
| CEN.PK113-7D | MATa MAL2-8c SUC2 | [58] |
| CB191 | CEN.PK113-7D *JEN1::GFP::KanMX* | This work |
| CB270 | CEN.PK113-7D *JEN1::GFP::HghMX* | This work |
| IMK302 | CEN.PK113-7D *jen1::loxP-KanMX4-loxP* | [59] |

### 4.2. Strain construction

All the *S. cerevisiae* strains generated in this work were constructed as follows: firstly, DNA fragments were amplified by PCR (Accuzyme DNA Polymerase, Bioline, or Supreme NZYProof DNA Polymerase, Nzytech) with specific oligonucleotides (listed in **Table 2**) using yeast genomic DNA or plasmid DNA. The resulting PCR products were introduced in *S. cerevisiae* cells, by following the polyethylene glycol (PEG)/lithium acetate (LiAc) protocol [60], and the resulting transformants were selected on geneticin (G418 disulfate, J62671) or hygromycin B (Alfa Aesar; J60681) according to the resistance (R) marker present (geneticin for *kan*R and hygromycin for *hgh*R).

**Table 2.**
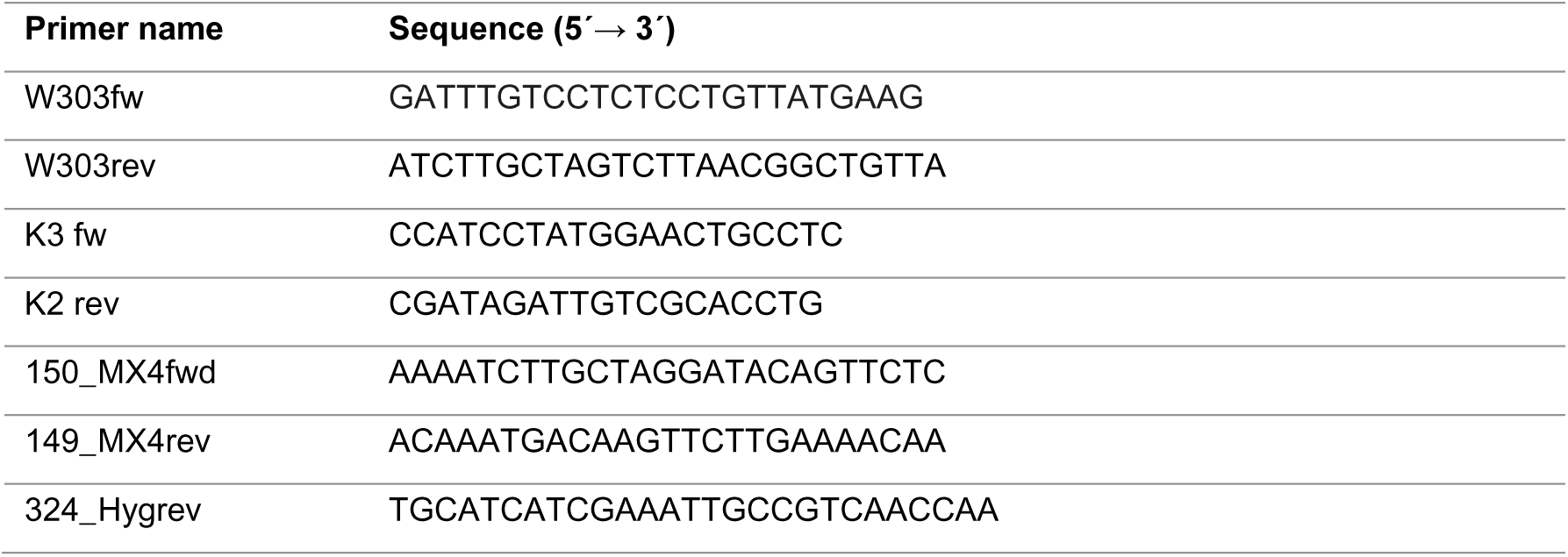
List of the primers used in this work.

Specifically, the strain CB191 (**Table 1**) was obtained as following: the DNA cassette *JEN1::GFP::kanMX* was PCR amplified from the genomic DNA of the strain Ace55 (**Table 1**) with the primers W303fw and W303rev. The DNA cassette was then introduced in the CENPK113-7D strain and positive transformants were confirmed by colony PCR using the primers K3 fw and W303rev or W303fw and K2 rev.

The CB270 strain (**Table 1**) was obtained by replacing the kanamycin resistance (*kanR) sequence* by the hygromycin resistance (*hghR)* DNA region in the CB191 strain. In detail, the *hghR* DNA sequence was first amplified from pAG32 plasmid [61] using primers 150_MX4fwd and 149_MX4rev (**Table 2**), which align 50 bps before the end of TEF promoter and 50 bps after the beginning of TEF terminator, respectively. Since KanR sequence is also flanked by TEF promoter and terminator regions, the 50 bp homology arms facilitate homologous recombination, allowing the replacement of *kanR* with *hghR*. Positive transformants were confirmed by colony PCR using primers 150 and 324.

### 4.3. Media and cultivation conditions

For the ‘quick’ and ethanol-only pH ramp experiments, *S. cerevisiae* pre-cultures were grown aerobically at 30 °C, 200 rpm, in 500 mL Erlenmeyer flasks containing 100 mL of synthetic medium (SM) [62] supplemented with 7.5 g/L of ethanol. Ethanol and filter-sterilised vitamins were added after autoclaving the other constituents of SM. For all other batch cultivations, 20 g/L of glucose was used instead of ethanol.

Continuous cultures were carried out in 2 L laboratory bioreactors (Getinge Applikon, Delft, The Netherlands) with a 1 L working volume. Cultures were grown aerobically at 30 °C, 800 rpm, in synthetic medium (for more details see [53, 62] and **Table S9**) with 33 mM (≈2.9 gL^−1^) L-lactic acid and 50 mM (≈2.3 gL^−1^) ethanol as sole carbon-limited sources. Ethanol and filter-sterilized vitamins (**Table S9** and [62]) were added after autoclaving the medium. After the batch phase, the medium pumps were switched on, resulting in the continuous addition of 70 mL/h of SM, supplemented with 33 mM L-lactic acid and 50 mM ethanol, to the culture. The working volume (1.0 L) was kept constant using an effluent pump controlled by an electric level sensor, resulting in a dilution rate (D) of 0.07 h^−1^. The culture pH was maintained at 6.0, 6.5, 7.0, 7.1 or 7.2 by automated addition of 2 M KOH or 2 M H_2_SO_4_. Chemostat cultures were considered to be in steady-state (SS) when, after at least 5 volume changes, the culture dry weight, extracellular metabolite concentrations, the CO_2_ production and O_2_ consumption rates did not differ more than 5 % over 2 volume changes. The chemostat runs were limited to less than 15-20 volume changes, after inoculation, to avoid evolutionary adaptation of the cultures.

For the pH ramp experiments, steady-state cultures were first established at pH 6.5 and subsequently a pH ramp from pH 6.5 to pH 7.5 was programmed in which the medium pH was automatically increased over the course of 120 hours (ethanol-only control) or 100 hours (Jen1-GFP, optical density was automatically monitored over the course of this experiment). For the ‘quick’ pH ramp experiments comparing WT to the otherwise congenic *Δjen1* strain, steady-state was established at pH 6.75, after which the pH ramp program increased the reactor pH setpoint 6.75 to pH setpoint 7.25 with 0.05 units per hour.

The off-gas signal (CO_2_ concentration) was automatically monitored over the course of all bioreactor experiments. Samples for dry weight, HPLC (for external metabolites analysis) and fluorescence microscopy (to follow the cellular localization of Jen1-GFP) were taken overtime.

The cultures were independently performed at least in duplicate (n≥2).

### 4.4. Analytical methods

#### 4.4.1. Optical density

Optical density at 660 nm was manually measured using a Jenway 7200 spectrophotometer (Bibby Scientific, Staffordshire, UK).

#### 4.4.2. Off-gas

Off-gas was first cooled in a condenser at 2 °C and then dried with a Perma Pure Dryer (Permapure, Toms River, NJ). Subsequently, CO_2_ and O_2_ concentrations were measured using an NGA 2000 Rosemount gas analyser (Emerson, St Louis, MO, USA).

#### 4.4.3. Dry weight

Culture dry weights were determined by filtrating 10 mL of culture using dry-, pre-weighed nitrocellulose filters with a pore size of 0.45 µm (Gelman Laboratory, Ann Arbor, MI, USA). The filters were washed twice with demineralized water before and after the filtration of the culture samples. After filtration, the filters were dried in a microwave for 20 min at 360 W and weighed again. Final dry weights are the average of duplicates performed for each chemostat culture.

### 4.5. High performance liquid chromatography (HPLC)

The extracellular metabolites and/or substrates present in the culture supernatants and media were analyzed by HPLC using an Agilent 1260 HPLC, equipped with a Bio-Rad HPX 87H ion-exchange column, operated at 60 °C with 5 mM H_2_SO_4_ as mobile phase at a flow rate of 0.600 mL·min^−1^. For sampling steady state cultures, ≈5 mL of culture broth from the bioreactor was rapidly sampled into a syringe containing pre-cooled stainless steel balls (4 mm, Fabory Laman Zoetermeer). The supernatant was immediately separated from the cells by filtration through a 0.45-µm pore size filter (PVD membrane, Merck Millipore Ltd.) ((adapted from [63]).

### 4.6. Epifluorescence microscopy and quantitative analysis

For the detection of Jen1-GFP cellular localization, a volume of 1 mL culture was collected and concentrated by a factor of 100 by centrifugation (5000 rpm,1 min). 4 μL of each sample was then directly and immediately visualized, without fixation, on a Zeiss Axio Imager Z1 microscope (Carl Zeiss AG, Oberkochen, Germany) with appropriate filters using a 100x magnification objective.

The quantitative analysis of the signal at PM/Total signal was performed using ImageJ software (version 1.53k) as described in [17, 64]. Specifically, two ellipses were drawn for each cell (n≥300) using the “magic wand” tool: the first ellipse was drawn around the entire cell (total fluorescence), and the second ellipse was drawn inside the cell, to exclude the plasma-membrane localized signal. The integrated density (IntDens) of both of these regions of interest was measured. The difference between the IntDens of the 1st ellipse (total cell) and the second ellipse (inside cell) gives the signal at the plasma membrane (signal at PM = IntDens[1st ellipse]−IntDens[2nd ellipse]). Data were represented as the ratio between the signal at PM and the total signal. An ordinary one-way ANOVA analysis was used followed by a Tukeýs multiple comparisons test using Prism 9.0 (GraphPad software, version 9.2.0). The *P* values are indicated (NS: *P*>0.05; *, *P*<0.05; **, *P*<0.01; ***, *P*<0.001; ****, *P*<0.0001).

### 4.7. RNA extraction, sequencing and analysis

#### 4.7.1. Culture sample collection

Steady state culture samples (± 240 mg of cells) were collected directly from the bioreactor and instantaneously frozen in liquid nitrogen and stored at – 80 °C for further RNA sample extraction.

#### 4.7.2. RNA extraction

For RNA extraction, ≈ 800 µL of “crushed ice” bits, from the – 80 °C samples, were transferred to Eppendorf tubes and thawed on ice. Eppendorfs were then centrifuged at 8000 *g*, 4 min, 0 °C and the pellet was resuspended in 200 µL cold RNase-free water. Subsequently, 400 µL of acid phenol-chloroform was quickly added and the mixture was centrifuged at 8000 *g*, for 4 min, at room temperature. The aqueous layer was transferred to a new tube and 400 µL of chloroform was added. The mixture was centrifuged again at 8000 *g*, 4 min, room temperature, and the upper layer was transferred to a new tube. 20 µL of 3 M Na-acetate and 500 µL of 100 % (v/v) ethanol (– 20 °C) were added. After mix by vortexing, the samples were incubated at – 20 °C for at least 15 min. The samples were then centrifuged for 15 min, 4°C, at 8000 *g* and the pellet was washed with 500 µL 80 % (v/v) ethanol (– 20 °C). The samples were vortexed, centrifuged again and, after removal of all the ethanol, the pellet was resuspended in 20 µL RNase-free water. The samples were kept at room temperature for at least 2 h to dissolve the RNA. The concentration, purity and quality of the total RNA samples were also determined using Tapestation (Agilent Technologies 2200) and NanoDrop 2000 (Thermo scientific).

#### 4.7.3. RNA sequencing

RNA samples [8 samples, four different pHs (6.0; 6.5; 7.0 and 7.1) in duplicate] were sent to Novogene (UK) for mRNA sequencing (RNA-seq). Non-directional libraries were prepared and checked with Qubit and real-time PCR for quantification and bioanalyzer for size distribution detection. Quantified libraries were pooled and sequenced on an Illumina platform.

#### 4.7.4. Analysis of RNA-Seq data

Raw reads (fastq format) were processed through FASTQ software [65] to yield clean data by removing low quality reads and reads containing adapters. The quality of the data was examined with the FastQC software (https://www.bioinformatics.babraham.ac.uk/projects/fastqc/). The average quality per read (Phred score) was above 35 for all samples. Mapping of paired-end clean reads (2×150 nt) were aligned to the *S. cerevisiae* CEN.PK113-7D genome (assembly ASM26988v1, obtained from https://fungi.ensembl.org/) with the Hisat2 v2.0.5 software [66]. Percentages of overall alignment were between 92 and 98%. The featureCounts (v1.5.0-p3) program was used to count the number of reads mapped to each gene. Differential expression analysis was conducted using DESeq2 R package (1.20.0) using the Benjamini and Hochberg’s approach to adjust the resulting *p*-values [67]. Genes with a *p*-value <=0.01 found by DESeq2 and a log_2_ fold change of +/− 0.6 were assigned as differentially expressed. Gene Ontology analyses for functional profiling were done with the g:Profiler server (https://biit.cs.ut.ee/ gprofiler/gost) using *S. cerevisiae* S288c strain data. The statistical domain scope used was “only annotated genes” and the g:SCS algorithm was applied for significance threshold (set at a *p*-value of 0.05). The search for transcription factors binding sites in the promoters of the selected genes was carried out at the Regulatory Sequence Analysis Tools (RSAT) server (https://rsat.france-bioinformatique.fr/fungi/) with the matrix-scan tool [68] and selecting an upstream segment of 800 nt (allowing overlapping with ORF). The transcription factor matrix data from the JASPAR database [69] was used and a *p*-value of 10^−4^ was set as significance threshold. Four individual sets of 174 promoters randomly extracted were also analysed and used as reference.

RNA-Seq data was deposited at *Repositório de Dados da Universidade do Minho* (dataRepositóriUM) under study https://doi.org/10.34622/datarepositorium/LWNKLQ.

## 5. Acknowledgments

We thank Charlotte Koster (TU Delft) for assistance with some of the laboratory experiments. We are also grateful to George Diallinas (University of Athens) for the critical reading of this manuscript.

## 8. Supporting Information sample

This article contains supporting information files:

- Supporting information 1 (word file)
- Supporting information 2 (excel file) sample

