## Supporting information 1 for "Jen1 transport and endocytosis in yeast reveal pH-responsive strategies to prevent metabolites loss"

Tables S1, S2, S3, S4, S5, S6, S7, S8, S9

Figures S1, S2, S3, S4

**Table S1 – Externally determined pH, biomass dry weight and extracellular metabolite analysis via HPLC obtained during the ‘Ethanol-only pH ramp’ experiment with *S. cerevisiae* CEN.PK113-7D.**

The concentration values [square brackets] are expressed in mg per L of the culture medium. The values are of 2 independent experiments (n=2). n.d. = not detected. The exact concentration values of ethanol inside the medium vessels were also measured by HPLC (bottom line).

| **Cultivation time (h)** | **External pH** | | **[Biomass**  **Dry weight]** | | **[Ethanol]** | | **[Acetate]** | | **[Pyruvate]** | |
| --- | --- | --- | --- | --- | --- | --- | --- | --- | --- | --- |
|  | **R5** | **R6** | **R5** | **R6** | **R5** | **R6** | **R5** | **R6** | **R5** | **R6** |
| 69.9 | **6.53** | **6.45** | 1270 | 1335 | n.d. | n.d. | n.d. | n.d. | n.d. | n.d. |
| 92.9 | **6.48** | **6.48** | 1340 | 1330 | n.d. | n.d. | n.d. | n.d. | n.d. | n.d. |
| 113.9 | **6.65** | **6.64** | 1345 | 1345 | n.d. | n.d. | n.d. | n.d. | 0.18 | 1.50 |
| 138.1 | **6.85** | **6.85** | 1325 | 1320 | n.d. | n.d. | n.d. | n.d. | 1.50 | 1.67 |
| 144.6 | **6.89** | **6.89** | 1320 | 1315 | n.d. | n.d. | n.d. | n.d. | 1.67 | 1.67 |
| 163.4 | **7.07** | **7.08** | 1360 | 1340 | n.d. | n.d. | 7 | 7 | 2.55 | 2.47 |
| 168.1 | **7.10** | **7.14** | 1340 | 1350 | n.d. | n.d. | 8 | 7 | 2.99 | 2.82 |
| 185.9 | **7.23** | **7.23** | 1075 | 1045 | 68 | 62 | 506 | 472 | 4.14 | 4.31 |
| 209.9 | **7.49** | **7.45** | 447.5 | 462.5 | 793 | 777 | 896 | 901 | 2.95 | 3.26 |
| **Medium vessel** | n.d. | n.d. | n.d. | n.d. | 2278 | 2293 | n.d. | n.d. | n.d. | n.d. |

**Table S2 – Externally determined pH, OD_660_, and extracellular metabolite analysis via HPLC obtained during the ‘quick pH ramp’ experiment with *S. cerevisiae* CEN.PK113-7D (WT).**

The concentration values [square brackets] are expressed in mg per L of the culture medium. The values are of 2 independent experiments (n=2). n.d. = not detected. In all samples the ethanol concentration was below the detection limit of the HPLC (<2mg/L). The exact concentrations inside the medium vessels were also measured by HPLC (bottom line).

| **Time (h)** | **External pH** | | **OD_660_** | | **[L-lactate]** | | **[Pyruvate]** | | **[Acetate]** | |
| --- | --- | --- | --- | --- | --- | --- | --- | --- | --- | --- |
|  | **R5** | **R6** | **R5** | **R6** | **R5** | **R6** | **R5** | **R6** | **R5** | **R6** |
| 0 | 6.73 | 6.74 | 18.60 | 18.55 | 31.86 | 31.86 | 7.04 | 7.93 | n.d. | n.d. |
| 1 | 6.78 | 6.8 | 17.85 | 18.73 | 35.29 | 32.72 | 7.93 | 8.81 | n.d. | n.d. |
| 2 | 6.80 | 6.82 | 17.85 | 18.30 | 37.86 | 37.86 | 8.81 | 9.69 | n.d. | n.d. |
| 3 | 6.84 | 6.86 | 17.00 | 17.50 | 42.15 | 41.30 | 9.69 | 10.57 | n.d. | n.d. |
| 4 | 6.89 | 6.92 | 17.00 | 17.75 | 46.44 | 47.30 | 10.57 | 12.33 | n.d. | n.d. |
| 5 | 6.94 | 6.96 | 17.50 | 17.55 | 51.59 | 53.30 | 12.33 | 14.09 | n.d. | n.d. |
| 6 | 7.01 | 7.02 | 17.60 | 16.85 | 58.45 | 59.31 | 14.09 | 17.61 | n.d. | n.d. |
| 7 | 7.07 | 7.07 | 17.90 | 18.50 | 67.03 | 69.60 | 16.73 | 22.02 | n.d. | n.d. |
| 8 | 7.09 | 7.10 | 17.80 | 18.20 | 78.18 | 82.47 | 21.13 | 29.06 | n.d. | n.d. |
| 9 | 7.13 | 7.14 | 16.95 | 17.50 | 93.62 | 109.9 | 27.30 | 37.87 | n.d. | 5.90 |
| 10 | 7.18 | 7.19 | 17.27 | 17.77 | 109.9 | 113.3 | 35.22 | 49.31 | n.d. | 28.93 |
| **Medium vessel** | n.d. | n.d. | **[Ethanol]**  2200 | **[Ethanol]**  2176 | 2589 | 2641 | 0.88 | 0.88 | n.d. | n.d. |

**Table S3 – Externally determined pH, OD_660_, and extracellular metabolite analysis via HPLC obtained during the ‘quick pH ramp’ experiment with *S. cerevisiae* IMK302 (*Δjen1*).**

The concentration values [square brackets] are expressed in mg per L of the culture medium. The values are of 2 independent experiments (n=2). n.d. = not detected. In all samples the ethanol concentration was below the detection limit of the HPLC (<2mg/L). The exact concentrations inside the medium vessels were also measured by HPLC (bottom line).

| **Time (h)** | **External pH** | | **OD_660_** | | **[L-lactate]** | | **[Pyruvate]** | | **[Acetate]** | |
| --- | --- | --- | --- | --- | --- | --- | --- | --- | --- | --- |
|  | **R3** | **R4** | **R3** | **R4** | **R3** | **R4** | **R3** | **R4** | **R3** | **R4** |
| 0 | 6.76 | 6.75 | 10.75 | 8.85 | 2493 | 2476 | 28.18 | 16.73 | n.d. | n.d. |
| 1 | 6.82 | 6.81 | 10.57 | 9.35 | 2500 | 2485 | 28.18 | 15.85 | n.d. | n.d. |
| 2 | 6.82 | 6.82 | 10.90 | 9.75 | 2494 | 2484 | 27.30 | 15.85 | n.d. | n.d. |
| 3 | 6.86 | 6.86 | 9.60 | 8.80 | 2500 | 2495 | 27.30 | 15.85 | n.d. | n.d. |
| 4 | 6.92 | 6.91 | 10.15 | 9.00 | 2513 | 2494 | 27.30 | 15.85 | n.d. | n.d. |
| 5 | 6.97 | 6.95 | 9.50 | 9.00 | 2522 | 2501 | 27.30 | 15.85 | n.d. | n.d. |
| 6 | 7.02 | 7.01 | 9.45 | 8.85 | 2512 | 2500 | 26.42 | 14.97 | n.d. | n.d. |
| 7 | 7.07 | 7.07 | 10.15 | 9.30 | 2530 | 2498 | 25.54 | 14.97 | n.d. | n.d. |
| 8 | 7.11 | 7.10 | 10.30 | 9.90 | 2516 | 2503 | 25.54 | 14.97 | n.d. | n.d. |
| 9 | 7.15 | 7.14 | 10.80 | 9.70 | 2538 | 2506 | 24.66 | 14.09 | n.d. | n.d. |
| 10 | 7.20 | 7.19 | 9.73 | 9.50 | 2532 | 2517 | 24.66 | 14.09 | 14.76 | 22.44 |
| **Medium vessel** | n.d. | n.d. | **[Ethanol]**  2192 | **[Ethanol]**  2002 | 2595 | 2633 | 0.88 | 0.88 | n.d. | n.d. |

Table S4 - Extracellular concentrations of ethanol and lactate obtained during the 'pH ramp' experiment with the CB270 Jen1-GFP strain.

The concentration values (C_out_) are expressed in mg of substrate (ethanol or lactate) per L of the culture medium (mg/L). The values are the mean of 2 independent experiments (n=2). s.d. and n.d. represent standard deviation and not detected, respectively. The exact concentration values of ethanol and lactate inside the medium vessels were also measured by HPLC (bottom line).

| **C_out_** (mg/L) | | | | |
| --- | --- | --- | --- | --- |
| **External pH** | **[Ethanol]** | | **[L-Lactate]** | |
|  | **Mean** | s.d. | **Mean** | s.d. |
| **6.50** | **n.d.** | n.d. | **13** | 1 |
| **6.75** | **n.d.** | n.d. | **6** | 1 |
| **7.00** | **n.d.** | n.d. | **9** | 1 |
| **7.25** | **n.d.** | n.d. | **52** | 23 |
| **7.50** | **356** | 72 | **494** | 101 |
| **Medium vessel** | **2317**  (≈2.3 g/L) | 0.014 | **2887**  (≈2.9 g/L) | 0.056 |

Table S5 - Dry weight values obtained at different extracellular pHs during the pH ramp experiment.

The values are the mean of two independent experiments (n=2) and s.d. represent standard deviation

| **External pH** | **6.50** | | **6.75** | | **7.00** | | **7.25** | | **7.50** | |
| --- | --- | --- | --- | --- | --- | --- | --- | --- | --- | --- |
|  | **Mean** | s.d. | **Mean** | s.d. | **Mean** | s.d. | **Mean** | s.d. | **Mean** | s.d. |
| **D.W.** (g/L) | **2.33** | 0.00 | **2.29** | 0.01 | **2.34** | 0.01 | **2.63** | 0.06 | **1.51** | 0.18 |

Table S6 - Extracellular concentrations of ethanol and lactate obtained for the steady state cultures set up at different pHs (indicated).

The concentration values are expressed in mg of substrate (lactate or ethanol) per L of the culture medium (left) or per L of the medium (right). The values are the mean of at least 2 independent experiments (n≥2). s.d. and n.d. represent standard deviation and not detected, respectively

| **C_out_** (mg/L) – from **culture medium** | | | | | **C_out_** (mg/L) – from **medium vessel** | | | |
| --- | --- | --- | --- | --- | --- | --- | --- | --- |
| **External pH** | **[Ethanol]** | | **[L-Lactate]** | | **[Ethanol]** | | **[L-Lactate]** | |
|  | **Mean** | s.d. | **Mean** | **s.d.** | **Mean** | s.d. | **Mean** | s.d. |
| **6.00** | **n.d.** | - | **10.795** | 6.795 | 2275 | 30 | 2834 | 19 |
| **6.50** | **n.d.** | - | **13.450** | 0.636 | 2317 | 13 | 2887 | 56 |
| **7.00** | **n.d.** | - | **33.800** | 3.629 | 2295 | 14 | 2804 | 28 |
| **7.10** | **n.d.** | - | **91.750** | 10.253 | 2294 | 16 | 2818 | 29 |

Table S7 - Extracellular concentrations of metabolites obtained for the steady state cultures set up at different pHs (indicated).

The concentration values are expressed in mg of substrate per L of the culture medium. The values are the mean of at least 2 independent experiments (n≥2). n.d. represent standard deviation and not detected, respectively

| **C_out_** (mg/L) | | | | | | | | | | |
| --- | --- | --- | --- | --- | --- | --- | --- | --- | --- | --- |
| **External pH** | **[Pyruvate]** | | **[Acetate]** | | **[Succinate]** | | **[Citrate]** | | **[Glycerol]** | |
|  | **Mean** | s.d. | **Mean** | s.d. | **Mean** | s.d. | **Mean** | s.d. | **Mean** | s.d. |
| **6.00** | **n.d.** | - | **30.400** | 13.859 | **15.450** | 7.283 | **2.285** | 0.219 | **n.d.** | - |
| **6.50** | **3.990** | 0.170 | **19.600** | 4.667 | **43.750** | 1.202 | **n.d.** | - | **n.d.** | - |
| **7.00** | **13.000** | 2.030 | **21.050** | 1.597 | **32.700** | 13.947 | **n.d.** | - | **6.253** | 2.910 |
| **7.10** | **92.750** | 20.153 | **198.500** | 98.288 | **51.800** | 15.556 | **n.d.** | - | **3.423** | 0.220 |

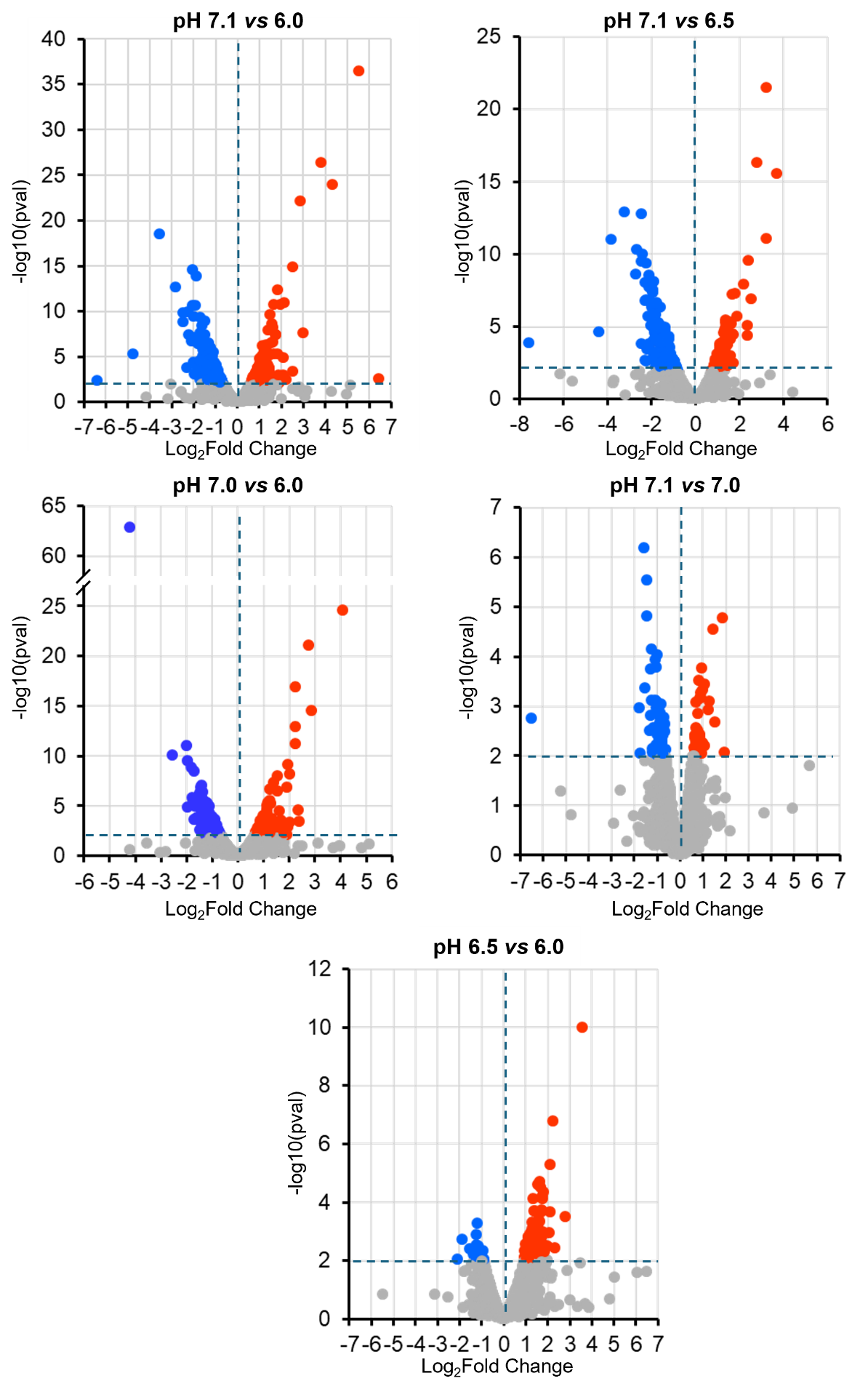

Fig. S1 - Vulcano plots of the different comparison made with the RNA-seq data presented in this work. Grey colour indicates values whose log2 Fold-Change is > -0.6 but <0.6, or their p-value is >0.01. Blue and red represent down- and up-regulated genes, respectively.

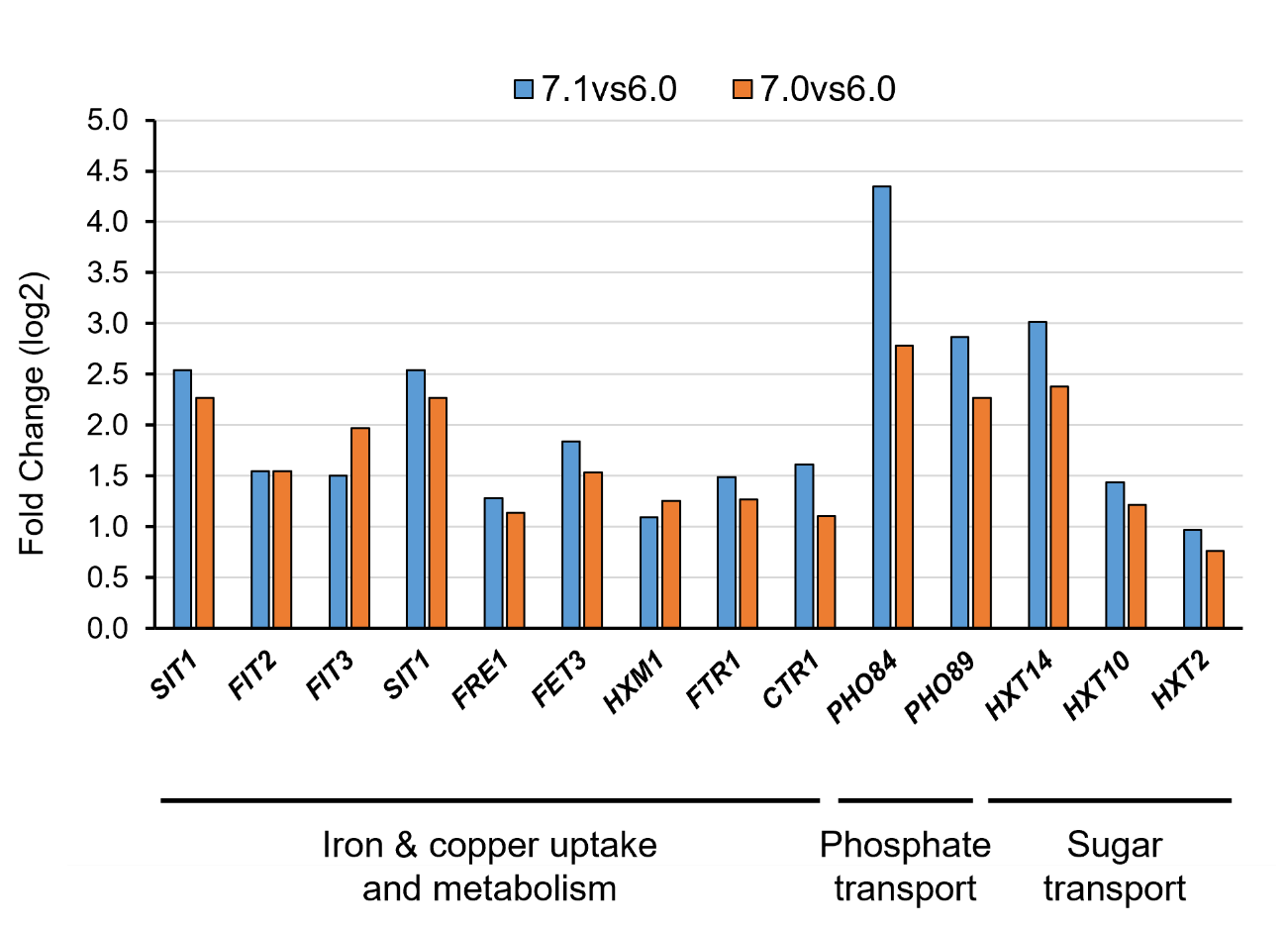

Fig. S2 - Changes in mRNA levels in cells shifted from pH 6.0 to pH 7.0 or 7.1 for a selection of genes typically induced by moderate alkalinization according to the literature (see Discussion)

**Table S8** - **Genes induced in the transition pH 6.0 to 7.1 but not when cells are shifted from pH 6.0 to 7.0.** Genes encoding proteins involved in mitochondrial translation are highlighted in red, those related to the electronic transport chain and ATP synthase function, are denoted in blue. Annotations combine GO-derived information and manual curation

| Gene_id | Standard Name | ORF | log2FoldChange | pvalue | Function Summary |
| --- | --- | --- | --- | --- | --- |
| CENPK1137D_4964 | LDS1 | YAL018C | 2.09 | 1.1E-03 | Protein involved in ascospore wall assembly; localizes to lipid droplets, prospore membrane, and ascospore wall |
| CENPK1137D_4994 | KTD1 | YAR028W | 1.51 | 8.9E-04 | Protein whose biological role is unknown; localized to the vacuole membrane in large scale studies |
| CENPK1137D_4907 | RFT1 | YBL020W | 0.81 | 5.0E-03 | Protein involved in glycolipid translocation; localizes to endoplasmic reticulum membrane |
| CENPK1137D_4893 | PET9 | YBL030C | 0.94 | 1.8E-04 | ATP/ADP exchanger of the mitochondrial inner membrane involved in respiration and apoptosis |
| CENPK1137D_4885 | MRPL16 | YBL038W | 0.68 | 9.5E-03 | Component of the large subunit of the mitochondrial ribosome, which mediates translation in the mitochondrion |
| CENPK1137D_4881 | ERD2 | YBL040C | 0.78 | 5.8E-03 | HDEL sequence-binding protein involved in protein retention in endoplasmic reticulum lumen and ER-to-Golgi vesicle-mediated transport; integral component of ER membrane |
| CENPK1137D_4863 | CMC2 | YBL059C-A | 1.11 | 9.2E-04 | Mitochondrial intermembrane space protein involved in respiratory chain complex assembly; also localizes to nucleus and cytoplasm |
| CENPK1137D_4916 | BNA4 | YBL098W | 1.01 | 3.6E-04 | Kynurenine 3-monooxygenase involved in de novo NAD biosynthesis from tryptophan; localizes to the mitochondrial outer membrane |
| CENPK1137D_4558 | GRX7 | YBR014C | 0.76 | 3.5E-03 | Glutathione-disulfide reductase involved in cellular response to oxidative stress; localizes to vacuole, golgi apparatus and lumen |
| CENPK1137D_4581 | SCO1 | YBR037C | 0.90 | 2.7E-03 | Putative thioredoxin peroxidase involved in protein complex assembly and copper ion transport; likely involved in response to redox state; localizes to mitochondrial inner membrane |
| CENPK1137D_4585 | FAT1 | YBR041W | 0.94 | 2.6E-03 | Long-chain fatty acid transporter involved in long-chain fatty acid uptake, protein quality control by the ubiquitin-proteasome system; localizes to ER, lipid droplet, peroxisome, cell periphery and vacuole membrane |
| CENPK1137D_4612 | BAP2 | YBR068C | 1.05 | 2.8E-04 | Amino acid transmembrane transporter that localizes to plasma membrane and endoplasmic reticulum |
| CENPK1137D_4694 | MRPS9 | YBR146W | 0.87 | 9.7E-04 | Component of the small subunit of the mitochondrial ribosome, which mediates translation in the mitochondrion |
| CENPK1137D_4711 | IFA38 | YBR159W | 0.76 | 3.0E-03 | Ketoreductase involved in biosynthesis of sphingolipids and very long-chain fatty acids; localizes to ER membrane |
| CENPK1137D_4737 | MBA1 | YBR185C | 1.31 | 9.8E-06 | Mitochondrial ribosome-binding protein involved in organization of the mitochondrial inner membrane; required for assembly of mitochondrial respiratory chain complexes III and IV; localizes to mitochondrial inner membrane |
| CENPK1137D_4760 | FTH1 | YBR207W | 0.67 | 6.2E-03 | Putative iron ion transporter involved in endocytosis; localizes to the vacuole membrane |
| CENPK1137D_4800 | ALG7 | YBR243C | 0.95 | 4.8E-04 | Dolichyl-P-dependent N-acetylglucosamine-1-P transferase (GPT) that catalyzes transfer of N-acetylglucosamine from UDP-N-acetylglucosamine to dolichyl phosphate; involved in synthesis of oligosaccharide precursor for N-linked protein glycosylation; localized in endoplasmic reticulum |
| CENPK1137D_4851 | VBA2 | YBR293W | 0.83 | 5.5E-03 | Basic amino acid transmembrane transporter localized to ER and vacuolar membranes |
| CENPK1137D_4519 | SLM5 | YCR024C | 1.01 | 1.3E-03 | Mitochondrial asparaginyl-tRNA synthetase that performs asparaginyl-tRNA aminoacylation by coupling asparagine to asparaginyl-tRNA |
| CENPK1137D_4530 | ELO2 | YCR034W | 0.83 | 8.8E-04 | Fatty acid elongase involved in very long-chain fatty acid biosynthesis, sphingolipid biosynthesis and late endosome to vacuole transport via the multivesicular body sorting pathway; localizes to the endoplasmic reticulum |
| CENPK1137D_4443 | GIT1 | YCR098C | 2.09 | 1.4E-05 | Glycerophosphodiester and glycerol-3-phosphate transmembrane transporter; involved in transport of glycerol-3-phosphate and glyceroldiester across plasma membrane |
| CENPK1137D_3820 | ATP16 | YDL004W | 0.68 | 5.9E-03 | Subunit of the central stalk of the F1 catalytic core of mitochondrial F1F0 ATP synthase; ATP synthase complex is localized to the mitochondrial inner membrane |
| CENPK1137D_3801 | MRX9 | YDL027C | 0.76 | 5.6E-03 | Protein whose biological role is unknown; localizes to the mitochondrion in a large-scale study |
| CENPK1137D_3784 | MTF2 | YDL044C | 1.01 | 1.0E-03 | RNA binding protein involved in mitochondrial translation and mRNA processing; localizes to mitochondria |
| CENPK1137D_3783 | FAD1 | YDL045C | 0.76 | 5.2E-03 | Cytoplasmic FMN adenylyltransferase involved in biosynthesis of flavin adenine dinucleotide (FAD) |
| CENPK1137D_3760 | IDP1 | YDL066W | 0.89 | 6.8E-04 | Mitochondrial isocitrate dehydrogenase (NADP+); has a role in glutamate biosynthesis |
| CENPK1137D_4253 | DLD2 | YDL178W | 0.76 | 2.4E-03 | Actin- and FAD-binding protein with D-2-hydroxyglutarate (D-2HG) dehydrogenase activity; oxidizes D-2HG to alpha-ketoglutarate; minor D-lactate dehydrogenase involved in lactate catabolism; localizes to mitochondrial matrix |
| CENPK1137D_4075 | GGC1 | YDL198C | 1.01 | 7.2E-05 | Guanine nucleotide transmembrane transporter involved in guanine nucleotide and transmembrane transport; localizes to mitochondria |
| CENPK1137D_3920 | SHR3 | YDL212W | 0.83 | 1.1E-03 | Protein chaperone of endoplasmic reticulum membrane, involved in COPII-coated vesicle budding; also localizes to mating projection tip |
| CENPK1137D_3860 | RSM10 | YDR041W | 1.02 | 9.7E-04 | Component of the small subunit of the mitochondrial ribosome, which mediates translation in the mitochondrion |
| CENPK1137D_3906 | TVP23 | YDR084C | 0.81 | 7.2E-03 | Golgi membrane protein involved in vesicle-mediated transport |
| CENPK1137D_3914 | ILT1 | YDR090C | 0.76 | 6.3E-03 | Predicted integral membrane protein whose biological role is unknown |
| CENPK1137D_3937 | MRPL1 | YDR116C | 0.98 | 5.5E-04 | Component of the large subunit of the mitochondrial ribosome, which mediates translation in the mitochondrion |
| CENPK1137D_3999 | RSM24 | YDR175C | 0.86 | 1.4E-03 | Component of the small subunit of the mitochondrial ribosome, which mediates translation in the mitochondrion |
| CENPK1137D_4018 | MSS116 | YDR194C | 1.12 | 2.0E-04 | RNA helicase involved in mRNA splicing, RNA folding, mitochondrial RNA processing and RNA elongation from mitochondrial promoter; localizes to mitochondrial matrix |
| CENPK1137D_4023 | CBS2 | YDR197W | 0.83 | 9.6E-03 | Mitochondrial translation regulator for the COB mRNA; localizes to mitochondrial inner membrane; associates with mitochondrial ribosomes |
| CENPK1137D_4093 | MSW1 | YDR268W | 1.06 | 3.9E-04 | Mitochondrial tryptophanyl-tRNA synthetase that performs tryptophanyl-tRNA aminoacylation by coupling tryptophan to tryptophanyl-tRNA |
| CENPK1137D_4121 | ATP5 | YDR298C | 0.69 | 5.2E-03 | Subunit of the stator stalk of the F0 coupling factor of mitochondrial F1F0 ATP synthase; ATP synthase complex is localized to the mitochondrial inner membrane |
| CENPK1137D_4150 | TIM11 | YDR322C-A | 0.90 | 2.0E-03 | Structural subunit of mitochondrial proton-transporting ATP synthase complex coupling factor F(o), involved in assembly of mitochondrial proton-transporting ATP synthase complex; also involved in cristae formation and ATP-synthesis coupled proton transport |
| CENPK1137D_4166 | MRPS28 | YDR337W | 0.84 | 7.9E-03 | Component of the small subunit of the mitochondrial ribosome, which mediates translation in the mitochondrion |
| CENPK1137D_2820 | HXT6 | YDR343C | 0.78 | 1.3E-03 | Hexose transmembrane transporter; also transports glucose, fructose, and mannose; localizes to mitochondria and plasma membrane |
| CENPK1137D_4203 | ARH1 | YDR376W | 0.73 | 8.5E-03 | Mitochondrial NADPH-adrenodoxin reductase involved in ubiquinone biosynthesis and iron homeostasis |
| CENPK1137D_4204 | ATP17 | YDR377W | 0.82 | 1.5E-03 | Subunit of the F0 coupling factor of mitochondrial F1F0 ATP synthase; ATP synthase complex is localized to the mitochondrial inner membrane |
| CENPK1137D_4222 | SHE9 | YDR393W | 1.26 | 2.5E-05 | Protein involved in organization of inner mitochondrial membrane |
| CENPK1137D_4346 | FPR2 | YDR519W | 0.71 | 6.1E-03 | Peptidylproyl isomerase that binds the macrolide lactone FK506; localizes to membrane, localizes to vacuoles, nucleus and cytoplasm in high-throughput studies |
| CENPK1137D_4357 | QCR7 | YDR529C | 0.83 | 3.7E-03 | Subunit of mitochondrial respiratory chain complex III, involved in electron transport and aerobic respiration |
| CENPK1137D_3651 | YEA4 | YEL004W | 0.93 | 2.8E-03 | UDP-N-acetylglucosamine transmembrane transporter involved in chitin biosynthesis; localizes to ER |
| CENPK1137D_3630 | RIP1 | YEL024W | 0.69 | 6.7E-03 | Ubiquinol-cytochrome-c reductase subunit of mitochondrial respiratory chain complex III; involved in aerobic respiration and mitochondrial electron transport |
| CENPK1137D_3616 | UTR2 | YEL040W | 1.18 | 6.1E-05 | Glycosyltransferase involved in chitin metabolism and cell wall organization; localizes to cell wall and bud neck septin ring |
| CENPK1137D_3594 | RML2 | YEL050C | 1.04 | 1.5E-04 | Component of the large subunit of the mitochondrial ribosome, which mediates translation in the mitochondrion |
| CENPK1137D_3572 | AFG1 | YEL052W | 0.71 | 9.3E-03 | Mitochondrial inner membrane protein involved in oxidative stress response and protein quality control |
| CENPK1137D_3673 | AFG3 | YER017C | 0.92 | 4.4E-04 | ATPase and metallopeptidase of the inner mitochondrial membrane; subunit of the m-AAA complex involved in assembly of mitochondrial membrane protein complexes, degradation of nonassembled inner membrane proteins and processing of proteins imported into mitochondria |
| CENPK1137D_3480 | VTC1 | YER072W | 0.85 | 7.6E-03 | mRNA-binding protein involved in vacuolar transport, fusion and microautophagy; also involved in polyphosphate metabolism; localizes to endoplasmic reticulum, vacuole membrane, and nuclear membrane |
| CENPK1137D_3531 | SHO1 | YER118C | 0.86 | 1.9E-03 | Membrane protein with osmosensor activity; subunit of the HICS complex; involved in cell polarity establishment and signaling for filamentous growth; localizes to the plasma membrane and bud |
| CENPK1137D_3367 | MDJ1 | YFL016C | 1.20 | 4.2E-04 | Unfolded protein-binding ATPase activator involved in heat response, protein folding and refolding, protein quality control, and mitochondrial genome maintenance; localizes to inner mitochondrial membrane |
| CENPK1137D_3350 | SNZ3 | YFL059W | 1.17 | 2.9E-03 | Pyridoxal 5'-phosphate synthase involved in pyridoxine and thiamine biosynthesis |
| CENPK1137D_3393 | MIC19 | YFR011C | 1.24 | 4.0E-05 | Subunit of mitochondrial MICOS complex involved in maintenance of crista junctions, inner membrane architecture, and formation of contact sites to outer mitochondrial membrane; localizes to mitochondrial crista junction |
| CENPK1137D_3402 | CSS2 | YFR020W | 1.30 | 1.3E-04 | Protein whose biological role is unknown; localizes to the endoplasmic reticulum and extracellular region |
| CENPK1137D_3426 | ERJ5 | YFR041C | 0.76 | 9.0E-03 | Endoplasmic reticulum protein involved in protein folding |
| CENPK1137D_3317 | KEG1 | YFR042W | 1.01 | 3.0E-03 | Membrane protein of the endoplasmic reticulum involved in 1,6-beta-glucan biosynthesis and chromosome organization |
| CENPK1137D_3320 | MRX20 | YFR045W | 0.75 | 9.5E-03 | Protein involved in mitochondrial transport; localizes to mitochondrial inner membrane |
| CENPK1137D_2960 | ERG4 | YGL012W | 0.71 | 3.8E-03 | Sterol reductase involved in ergosterol biosynthesis; localizes to endoplasmic reticulum |
| CENPK1137D_2891 | GUP1 | YGL084C | 0.69 | 9.5E-03 | O-acyltransferase involved in glycerol transport and catabolism, and GPI anchor biosynthesis; localizes to plasma membrane and endoplasmic reticulum |
| CENPK1137D_2867 | RMD9 | YGL107C | 0.87 | 2.0E-03 | Protein that binds mRNA 3'-UTR; involved in mitochondrial mRNA processing, and in regulation of mitochondrial transcription and translational initiation; localized to mitochondrial inner membrane |
| CENPK1137D_2831 | MRF1 | YGL143C | 0.98 | 3.0E-03 | Translation release factor involved in mitochondrial translational termination; localizes to mitochondria |
| CENPK1137D_2828 | RRT6 | YGL146C | 1.07 | 9.8E-04 | Predicted membrane protein whose biological role is unknown |
| CENPK1137D_3263 | EMP24 | YGL200C | 0.79 | 4.0E-03 | Protein involved in ER to Golgi vesicle-mediated transport, protein retention in ER lumen snd vesicle organization; localizes to COPII vesicle and ER |
| CENPK1137D_2983 | MCY1 | YGR012W | 0.73 | 6.2E-03 | Protein whose biological role is unknown; localizes to the mitochondrion and mitochondrial outer membrane in different large-scale studies |
| CENPK1137D_2996 | MSP1 | YGR028W | 0.79 | 6.8E-03 | ATPase involved in protein-mitochondrial targeting; localizes to peroxisomal and mitochondrial outer membrane |
| CENPK1137D_3031 | VHT1 | YGR065C | 1.25 | 1.6E-06 | Biotin transmembrane transporter of the plasma membrane |
| CENPK1137D_3074 | VOA1 | YGR106C | 0.79 | 2.9E-03 | Protein involved in cellular protein complex assembly; localizes to endoplasmic reticulum membrane and vacuole membrane |
| CENPK1137D_3081 | SHY1 | YGR112W | 0.96 | 1.9E-03 | Protein involved in mitochondrial respiratory chain complex IV assembly; integral component of the mitochondrial inner membrane |
| CENPK1137D_3089 | YGR122W | YGR122W | 0.86 | 9.3E-03 | Protein that may be involved in pH regulation |
| CENPK1137D_3129 | MRPS35 | YGR165W | 0.93 | 1.6E-03 | Component of the small subunit of the mitochondrial ribosome, which mediates translation in the mitochondrion |
| CENPK1137D_3153 | CRH1 | YGR189C | 0.75 | 1.6E-03 | Glucosyltransferase that cross-links chitin to beta(1-6) and beta(1-3) glucans in the cell wall; involved in cell wall organization; localized to sites of cell wall growth, such as incipient buds |
| CENPK1137D_3154 | HIP1 | YGR191W | 0.81 | 2.2E-03 | High-affinity L-histidine transmembrane transporter involved in histidine and manganese transport; localizes to cell periphery, plasma membrane, and endoplasmic reticulum |
| CENPK1137D_3173 | CIR1 | YGR207C | 1.00 | 6.5E-04 | Protein whose biological role is unknown; localizes to the mitochondrion in different large-scale studies |
| CENPK1137D_3182 | RSM27 | YGR215W | 0.99 | 1.9E-03 | Component of the small subunit of the mitochondrial ribosome, which mediates translation in the mitochondrion |
| CENPK1137D_3186 | MRPL9 | YGR220C | 1.00 | 1.2E-04 | Component of the large subunit of the mitochondrial ribosome, which mediates translation in the mitochondrion |
| CENPK1137D_3199 | YHB1 | YGR234W | 0.89 | 1.5E-03 | Nitric oxide reductase involved in response to reactive nitrogen species, oxidative stress, and misfolded protein; localizes to nucleus, mitochondria, and stress granules |
| CENPK1137D_3314 | MAL12 | YGR292W | 1.05 | 6.8E-03 | Protein with glucan 1,4-alpha-maltotriohydrolase, maltose alpha-glucosidase, and sucrose alpha-glucosidase activities; involved in sucrose and maltose catabolism |
| CENPK1137D_5387 | LAG1 | YHL003C | 0.95 | 4.2E-04 | Sphingosine N-acyltransferase subunit of the acyl-CoA ceramide synthase complex, involved in ceramide biosynthesis and replicative cell aging |
| CENPK1137D_5379 | YHL009W-A | YHL009W-A | 6.46 | 2.8E-03 | Retrotransposon TYA Gag gene co-transcribed with TYB Pol |
| CENPK1137D_5372 | YLF2 | YHL014C | 1.21 | 3.4E-04 | Protein of unknown function |
| CENPK1137D_5384 | ARN1 | YHL040C | 1.16 | 1.1E-05 | Siderophore transmembrane transporter that locailzes to cell periphery, cytoplasmic vesicles, endosome, vacuole, and nucleus |
| CENPK1137D_5449 | COX6 | YHR051W | 0.89 | 4.5E-04 | Subunit of the mitochondrial respiratory chain complex IV; contributes to cytochrome c oxidase activity and electron transport to oxygen |
| CENPK1137D_5237 | MSR1 | YHR091C | 0.88 | 4.6E-03 | Putative mitochondrial arginyl-tRNA synthetase that performs arginyl-tRNA aminoacylation by coupling arginine to arginyl-tRNA |
| CENPK1137D_5332 | SVP26 | YHR181W | 0.89 | 1.3E-03 | COPII receptor involved in ER to Golgi vesicle-mediated transport, cell wall organization and protein retention in Golgi apparatus; localizes to ER |
| CENPK1137D_5356 | SMN1 | YHR202W | 0.82 | 5.7E-03 | Protein whose biological role is unknown; localizes to the cytosol in a classical study and to the vacuole in a large scale study |
| CENPK1137D_5024 | PRM2 | YIL037C | 1.20 | 3.9E-03 | Membrane protein involved in cell fusion and karyogamy |
| CENPK1137D_5140 | RHO3 | YIL118W | 0.88 | 6.5E-03 | Cytosolic GTPase involved in activation of exocytosis, upregulation of actin cable assembly, and actin cytoskeleton polarity; localizes to plasma membrane and bud |
| CENPK1137D_5116 | AXL2 | YIL140W | 0.68 | 9.9E-03 | Protein involved in axial cellular bud site selection; localizes to bud neck and bud neck septin ring |
| CENPK1137D_1290 | SYS1 | YJL004C | 0.74 | 7.6E-03 | Protein involved in Golgi to endosome transport and protein localization to Golgi apparatus; localizes to trans-Golgi network |
| CENPK1137D_1475 | CIS3 | YJL158C | 0.67 | 5.1E-03 | Structural protein of the cell wall |
| CENPK1137D_1207 | NUC1 | YJL208C | 1.45 | 2.0E-04 | Endodeoxyribonuclease involved in apoptotic DNA fragmentation, DNA and RNA catabolism; localizes to cytosol, nucleus and mitochondria |
| CENPK1137D_1339 | CYC1 | YJR048W | 1.17 | 7.0E-05 | Electron carrier of the mitochondrial inter membrane space, involved in ubiquinol-to-cytochrome c and cytochrome c-to-oxygen electron transport |
| CENPK1137D_1370 | MIR1 | YJR077C | 0.83 | 5.4E-04 | Inorganic phosphate transmembrane transporter involved in phosphate ion transmembrane transport; integral mitochondrial inner membrane; localizes to mitochondria |
| CENPK1137D_1372 | AIM24 | YJR080C | 1.18 | 6.0E-06 | Protein with a role in organization of the mitochondrial inner membrane; integral to the inner membrane |
| CENPK1137D_1394 | RSM26 | YJR101W | 0.99 | 5.9E-04 | Component of the small subunit of the mitochondrial ribosome, which mediates translation in the mitochondrion |
| CENPK1137D_1415 | ATP2 | YJR121W | 0.94 | 1.6E-04 | Subunit of the catalytic core of the F1 sector of mitochondrial F1F0 ATP synthase; ATP synthase complex is localized to the mitochondrial inner membrane |
| CENPK1137D_996 | CCE1 | YKL011C | 1.06 | 3.4E-03 | Endodeoxyribonuclease of the mitochondrial inner membrane; involved in mitochondrial genome maintenance |
| CENPK1137D_920 | HOT13 | YKL084W | 0.94 | 5.6E-03 | Zinc-binding protein involved in mitochondrial protein import; localizes to mitochondrial intermembrane space |
| CENPK1137D_917 | CYT2 | YKL087C | 1.87 | 1.1E-03 | Holocytochrome c synthase involved in cytochrome c-heme linkage; localizes to mitochondrial intermembrane space |
| CENPK1137D_1188 | VPH2 | YKL119C | 0.93 | 3.3E-03 | Subunit of Vma12-Vma22 assembly complex, involved in vacuolar acidification and assembly of the V-type ATPase complex; localizes to ER membrane |
| CENPK1137D_1173 | OCT1 | YKL134C | 0.84 | 4.7E-03 | Mitochondrial metalloendopeptidase involved in protein processing and targeting, protein stabilization, and iron homeostasis |
| CENPK1137D_1171 | CMC1 | YKL137W | 0.89 | 4.8E-03 | Copper-binding protein of the mitochondrial intermembrane space; involved in assembly of cytochrome c oxidase |
| CENPK1137D_1166 | SDH3 | YKL141W | 0.72 | 8.6E-03 | Subunit of the succinate dehydrogenase mitochondrial respiratory chain complex, and also of the TIM22 mitochondrial inner membrane protein insertion complex; required for cellular respiration |
| CENPK1137D_1145 | MCD4 | YKL165C | 0.76 | 7.6E-03 | Protein with mannose-ethanolamine phosphotransferase activity; involved in ATP transport and in biosynthesis of glycosylphosphatidylinositol (GPI) anchors; localizes to the vacuole, cell wall, and endoplasmic reticulum |
| CENPK1137D_1080 | MST1 | YKL194C | 1.27 | 2.1E-04 | Mitochondrial threonyl-tRNA synthetase that performs threonyl-tRNA aminoacylation in the mitochondria by coupling threonine to threonyl-tRNA |
| CENPK1137D_1152 | SRY1 | YKL218C | 0.88 | 2.0E-03 | Threo-3-hydroxyaspartate ammonia-lyase involved in catabolism of modified amino acids |
| CENPK1137D_1022 | PRY2 | YKR013W | 0.93 | 3.2E-04 | Sterol binding protein; involved in the export of free fatty acids and sterols; localizes to the extracellular region |
| CENPK1137D_1026 | MIC60 | YKR016W | 1.09 | 5.4E-04 | Subunit of MICOS, a mitochondrial inner membrane complex; involved in mitochondrial cristae formation and the import of proteins into the mitochondrial intermembrane space |
| CENPK1137D_1041 | GMH1 | YKR030W | 1.20 | 2.5E-05 | Golgi integral membrane protein involved in vesicle-mediated transport |
| CENPK1137D_1099 | OMA1 | YKR087C | 0.74 | 9.1E-03 | Metalloendopeptidase involved in mitochondrial protein catabolism, response to redox state and TOR signaling; localizes to mitochondrial inner membrane |
| CENPK1137D_455 | MEF1 | YLR069C | 1.06 | 7.5E-04 | Mitochondrial translation elongation factor |
| CENPK1137D_570 | PEX13 | YLR191W | 0.89 | 3.5E-03 | Subunit of the peroxisomal importomer complex involved in the docking of proteins in the peroxisomal matrix |
| CENPK1137D_614 | LIP2 | YLR239C | 0.95 | 2.3E-03 | Mitochondrial ligase involved in protein lipoylation |
| CENPK1137D_625 | SSP120 | YLR250W | 0.93 | 2.3E-03 | Protein whose biological role is unknown; localizes to the cytoplasm in a large-scale study |
| CENPK1137D_627 | CQD2 | YLR253W | 0.99 | 5.7E-04 | Protein involved in mitochondrion organization during lipid homeostatis; is an integral component of the mitochondrial inner membrane |
| CENPK1137D_686 | MRPL15 | YLR312W-A | 0.98 | 2.6E-03 | Component of the large subunit of the mitochondrial ribosome, which mediates translation in the mitochondrion |
| CENPK1137D_698 | YLR326W | YLR326W | 0.93 | 9.3E-04 | Protein whose biological role is unknown; localizes to the cell periphery |
| CENPK1137D_702 | JIP3 | YLR331C | 1.35 | 1.4E-04 | Protein whose biological role and cellular location are unknown |
| CENPK1137D_742 | ELO3 | YLR372W | 0.72 | 3.4E-03 | Fatty acid elongase involved in very long-chain fatty acid biosynthesis, sphingolipid biosynthesis and late endosome to vacuole transport via the multivesicular body sorting pathway; localizes to the endoplasmic reticulum |
| CENPK1137D_54 | MRPL3 | YMR024W | 1.21 | 1.3E-05 | Component of the large subunit of the mitochondrial ribosome, which mediates translation in the mitochondrion |
| CENPK1137D_166 | ERG29 | YMR134W | 0.84 | 2.6E-03 | Protein involved in iron homeostasis, ergosterol biosynthesis, and mitochondrion organization; localizes to cytoplasm, endoplasmic reticulum, and nuclear envelope |
| CENPK1137D_177 | NDE1 | YMR145C | 0.73 | 1.7E-03 | NADH dehydrogenase involved glycolytic fermentation to ethanol and NADH oxidation; also involved in regulation of apoptosis; localizes to mitochondria |
| CENPK1137D_192 | ATG16 | YMR159C | 2.23 | 3.7E-03 | Atg8 ligase (conjugates Atg8p to phosphatidylethanolamine, PE); subunit of the Atg12-Atg5-Atg16 complex; involved in macroautophagy, cytoplasm-to-vacuole targeting (CVT) pathway, late nucleophagy, piecemeal microautophagy of nucleus, protein lipidation and mitochondrion degradation; localizes to the pre-autophagosomal structure (PAS) |
| CENPK1137D_222 | MRPS17 | YMR188C | 0.96 | 7.4E-03 | Component of the small subunit of the mitochondrial ribosome, which mediates translation in the mitochondrion |
| CENPK1137D_267 | MTF1 | YMR228W | 0.89 | 2.3E-03 | Mitochondrial transcription factor involved in transcription initiation from mitochondrial promoter and activation of RNA elongation; subunit of mitochondrial DNA-directed RNA polymerase complex; localizes to mitochondrial matrix and intermembrane space |
| CENPK1137D_348 | YME2 | YMR302C | 0.67 | 9.4E-03 | Mitochondrial inner membrane protein involved in mitochondrial genome maintenance |
| CENPK1137D_350 | SCW10 | YMR305C | 0.77 | 3.0E-03 | Cell wall protein with possible glucosidase activity; appears to be involved in conjugation |
| CENPK1137D_352 | GAS1 | YMR307W | 0.69 | 3.7E-03 | 1,3-beta-glucanosyltransferase, catalyzes splitting and linking of (1->3)-beta-D-glucan molecules that leads to elongation of (1->3)-beta-D-glucan chains; involved in cell wall organization, also has a role in chromatin silencing; localized to cell wall, nuclear periphery and throughout secretory pathway |
| CENPK1137D_2671 | MRP7 | YNL005C | 0.77 | 5.0E-03 | Component of the large subunit of the mitochondrial ribosome, which mediates translation in the mitochondrion; may have a role in ribosomal peptidyltransferase activity |
| CENPK1137D_2578 | MIC27 | YNL100W | 0.82 | 2.4E-03 | Protein involved in cristae formation and protein-containing complex subunit organization; subunit of MICOS complex; localizes to vacuole membrane and mitochondria |
| CENPK1137D_2559 | TOM70 | YNL121C | 0.75 | 6.6E-03 | Protein transmembrane transporter with mitochondrion targeting sequence binding activity involved in protein targeting to mitochondrion, protein import into mitochondrial matrix, and in protein insertion into mitochondrial inner membrane; subunit of mitochondrial outer membrane translocase complex |
| CENPK1137D_2548 | TOM22 | YNL131W | 0.68 | 8.1E-03 | Protein transmembrane transporter involved in protein import into mitochondrial matrix and protein insertion into mitochondrial outer membrane; subunit of mitochondrial outer membrane translocase complex |
| CENPK1137D_2542 | NAM9 | YNL137C | 1.13 | 4.9E-05 | Component of the small subunit of the mitochondrial ribosome, which mediates translation in the mitochondrion |
| CENPK1137D_2756 | MRPL10 | YNL284C | 0.89 | 1.1E-03 | Component of the large subunit of the mitochondrial ribosome, which mediates translation in the mitochondrion |
| CENPK1137D_2774 | PFA3 | YNL326C | 0.90 | 3.0E-03 | Vacuolar membrane palmitoyltransferase involved in non-autophagic vacuole fusion |
| CENPK1137D_2014 | CSI2 | YOL007C | 1.10 | 1.3E-03 | Protein whose biological role is unknown; localizes to the bud neck and vacuole |
| CENPK1137D_2000 | TAT2 | YOL020W | 0.78 | 7.7E-03 | Aromatic amino acid transmembrane transporter involved in tryptophan transport; localizes to eisosome, plasma membrane and ER |
| CENPK1137D_1997 | IFM1 | YOL023W | 1.03 | 6.4E-03 | Mitochondrial translation initiation factor; binds tRNA and has GTPase activity |
| CENPK1137D_1992 | MDM38 | YOL027C | 0.69 | 7.5E-03 | Ribosome binding protein that positively regulates mitochondrial translation with Mba1p; involved in cellular potassium ion homeostasis by transporting potassium ions with Rbk1p and YDL183Cp; plays a role in biogenesis of both cytochrome bc(1) and cytochrome c oxidase complexes; localized to mitochondrial inner membrane |
| CENPK1137D_1967 | AIM39 | YOL053W | 0.80 | 3.6E-03 | Protein whose biological role is unknown; localizes to the mitochondrion in a large scale study |
| CENPK1137D_2414 | ITR2 | YOL103W | 0.92 | 1.8E-03 | Myo-inositol transmembrane transporter that localizes to cell periphery, plasma membrane, and vacuole |
| CENPK1137D_1982 | FRE7 | YOL152W | 1.35 | 6.5E-05 | Iron chelate reductase involved in copper and iron transport; localizes to plasma membrane |
| CENPK1137D_2085 | CYT1 | YOR065W | 0.96 | 4.8E-04 | Subunit of mitochonrial respiratory complex III, acts as an electron carrier that moves electrons within the ubiquinol-cytochrome c reductase complex; localized to mitochondrial inner membrane |
| CENPK1137D_2098 | ATX2 | YOR079C | 0.85 | 4.7E-03 | Manganese ion transmembrane transporter involved in manganese ion homeostasis; localizes to endoplasmic reticulum, Golgi membrane, trans-Golgi network, and late endosome |
| CENPK1137D_2122 | PIN2 | YOR104W | 0.95 | 2.6E-03 | Predicted integral membrane protein whose biological role is unknown |
| CENPK1137D_2153 | IDH2 | YOR136W | 0.77 | 9.3E-03 | Catalytic subunit of the mitochondrial isocitrate dehydrogenase complex; has roles in isocitrate metabolism, glutamate biosynthesis, and the tricarboxylic acid cycle |
| CENPK1137D_2203 | TUF1 | YOR187W | 1.13 | 3.4E-05 | Mitochondrial translation elongation factor, catalyzes the binding of aminoacyl-tRNA to ribosomes and possesses an intrinsic GTPase activity |
| CENPK1137D_2231 | SPR2 | YOR214C | 2.52 | 4.1E-04 | Protein whose biological role is unknown; localizes to the cell wall |
| CENPK1137D_2303 | MPD1 | YOR288C | 0.84 | 3.6E-03 | Protein disulfide isomerase involved in protein folding; localizes to ER and vacuole |
| CENPK1137D_2323 | RRG7 | YOR305W | 1.15 | 8.7E-03 | Protein whose biological role is unknown; localizes to the mitochondrion in a large-scale study |
| CENPK1137D_1661 | ISM1 | YPL040C | 1.09 | 1.6E-03 | Mitochondrial isoleucine-tRNA ligase that couples isoleucine to isoleucyl-tRNA |
| CENPK1137D_1651 | MNN9 | YPL050C | 0.67 | 9.1E-03 | Alpha-1,6-mannosyltransferase involved in protein N-linked glycosylation; localizes to ER and cis-Golgi network |
| CENPK1137D_1643 | SUR1 | YPL057C | 0.68 | 6.4E-03 | Catalytic subunit of mannosylinositol phosphorylceramide (MIPC) synthase; required for biosynthesis of mature sphingolipids; predicted to be integral to membranes |
| CENPK1137D_1621 | ATP4 | YPL078C | 0.75 | 2.3E-03 | Subunit of stator stalk of F0 coupling factor of mitochondrial F1F0 ATP synthase; also has a role in oligomerization of the ATP synthase complex, which is localized to mitochondrial inner membrane; involved in ATP-synthesis coupled proton transport |
| CENPK1137D_1613 | YDC1 | YPL087W | 0.79 | 5.0E-03 | Dihydroceramidase involved in ceramide metabolism; localizes to endoplasmic reticulum and vacuole membrane |
| CENPK1137D_1603 | MSY1 | YPL097W | 1.23 | 1.2E-04 | Mitochondrial tyrosyl-tRNA synthetase that performs mitochondrial tyrosyl-tRNA aminoacylation by coupling tyrosine to tyrosyl-tRNA |
| CENPK1137D_1597 | FMP30 | YPL103C | 0.82 | 8.7E-03 | Putative NAPE-specific phospholipase D involved in N-acylethanolamine and N-acylphosphatidylethanolamine metabolism; localizes to mitochondria |
| CENPK1137D_1596 | MSD1 | YPL104W | 0.98 | 6.7E-04 | Mitochondrial aspartyl-tRNA synthetase that performs aspartyl-tRNA aminoacylation by coupling aspartate to aspartyl-tRNA |
| CENPK1137D_1566 | COX11 | YPL132W | 0.96 | 8.7E-04 | Mitochondrial inner membrane protein that binds copper and is involved in respiratory chain complex assembly |
| CENPK1137D_1526 | COX10 | YPL172C | 1.10 | 1.7E-03 | Putative protoheme IX farnesyltransferase, catalyzes conversion of protoheme to heme A in heme A biosynthesis; localized to mitochondria in high-throughput experiments |
| CENPK1137D_1918 | TYW1 | YPL207W | 0.81 | 6.2E-03 | S-adenosyl-L-methionine binding protein involved in wybutosine biosynthesis; localizes to endoplasmic reticulum |
| CENPK1137D_1855 | VMA11 | YPL234C | 0.83 | 2.4E-03 | Subunit of V0 vacuolar domain of V-ATPase involved in vacuolar acidification; localizes to vacuole membrane and cell periphery |
| CENPK1137D_1733 | RBD2 | YPL246C | 0.75 | 8.3E-03 | Predicted integral membrane protein whose biological role is unknown; colocalizes with COPI-coated vesicles and localizes to the Golgi apparatus in multiple large-scale studies; localizes to the nuclear periphery |
| CENPK1137D_1524 | SAM3 | YPL274W | 0.68 | 5.4E-03 | Plasma membrane protein involved in transporting spermidine, putrescine, and S-adenosyl-L-methionine |
| CENPK1137D_1726 | YME1 | YPR024W | 0.83 | 2.3E-03 | ATP-dependent peptidase of the inner mitochondrial membrane; subunit of the i-AAA complex involved in mitochondrial protein turnover; also involved in protein import, folding and maturation |
| CENPK1137D_1728 | YPR027C | YPR027C | 0.91 | 8.1E-03 | Protein whose biological role is unknown; localizes to the endoplasmic reticulum |
| CENPK1137D_1749 | MSF1 | YPR047W | 0.91 | 1.2E-03 | Mitochondrial phenylalanyl-tRNA synthetase that catalyzes phenylalanyl-tRNA aminoacylation through the coupling of phenylalanine to phenylalanyl-tRNA |
| CENPK1137D_1780 | DIB1 | YPR082C | 0.91 | 5.2E-03 | Subunit of the U4/U6 x U5 tri-small nuclear RNP (snRNP) complex involved in messenger RNA (mRNA) splicing via spliceosome; localizes to U5 snRNP |
| CENPK1137D_1831 | LOA1 | YPR139C | 1.05 | 6.8E-05 | Lysophosphatidic acid acyltransferase that transfers acyl groups from an acyl-CoA to lysophosphatidic acid to form phosphatidic acid; involved in triglyceride homeostasis and lipid droplet organization; localized to the ER and lipid droplets |
| CENPK1137D_1850 | KRE6 | YPR159W | 0.71 | 7.0E-03 | Glucosyl hydrolase (transglucosidase); involved in biosynthesis of Î² (1->6)-D-glucan, a key component of the cell wall; integral component of membranes of the ER and transport vesicles; and of the plasma membrane |

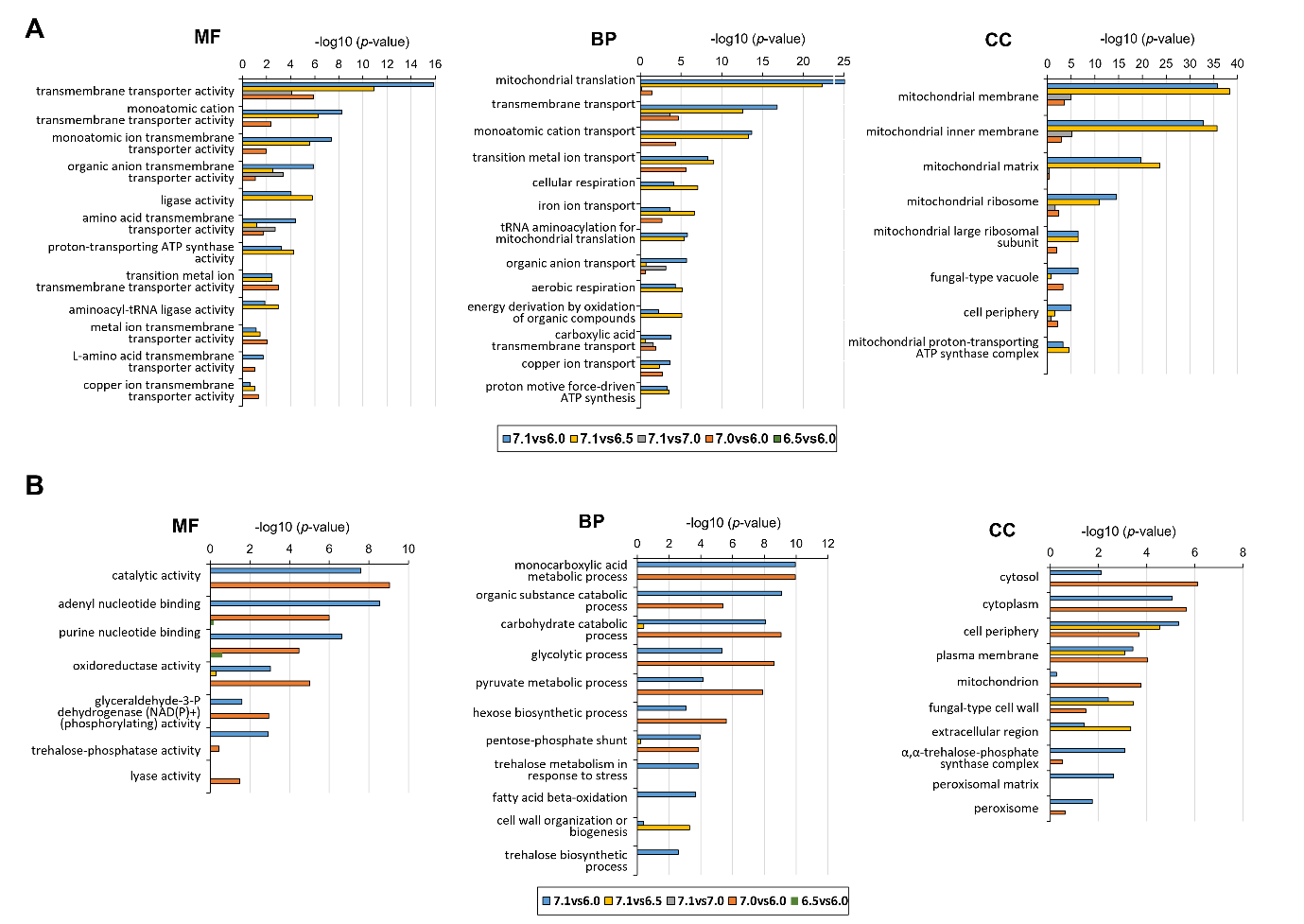

Fig. S3 – Multiple comparisons of enrichment for specific Gene Ontology terms of differentially up-regulated (A) and down-regulated (B) genes. MF, molecular function; BP, biological process; CC, cellular component. The pH comparisons are represented by colour bars: blue, for pH7.1vspH6.0; yellow, for pH7.1vspH6.5; grey, for pH7.1vspH7.0; orange, for pH7.0vspH6.0 and green, for pH6.5vspH6.0.The p-value threshold for significance was set at 0.05.

Table S9 - Detailed composition of the synthetic medium (SM) for carbon limited-, aerobic- continuous culture of *S. cerevisiae* (final volume: 20 L)

| Component | Formula | Product code | MW  (g.mol^-1^) | Weight  (g) | Volume (mL) | Comment |
| --- | --- | --- | --- | --- | --- | --- |
| Ammonium Sulphate | (NH_4_)_2_SO_4_ | Merck 1211 | 132.14 | 100.00 | - | - |
| Potassium dihydrogen phosphate | KH_2_PO_4_ | Merck  4877 | 136.09 | 60.00 | - | - |
| Magnesium sulphate. 7H_2_O | MgSO_4_.7H_2_O | Baker 0168 | 246.47 | 10.00 | - | - |
| Pluronic PE6100 | - | BASF 50073570 | - | 8.00 | - | - |
| Trace metals solution | - | *Stock solution | - | - | 20.00 | - |
| L-(+)-Lactic acid | C_3_H_6_O_3_ | - | 90.08 | - | 61.93 | - |
| RO water (total weight) | H_2_O | - | - | 19422 | - | - |
| Ethanol | CH_3_CH_2_OH | - | 46.07 | - | 58.20 | Ethanol and vitamins were added after autoclaving the above components |
| Vitamins | - | **Stock solution | - | - | 20.00 |  |

*The **trace metals stock solution** contained (L^-1^):

- 4.5 g CaCl_2_.2H_2_O
- 4.5 g ZnSO_4_.7H_2_O
- 3 g FeSO_4_.7H_2_O,
- 1 g H_3_BO_3_,
- 1 g MnCl_2_.4H_2_O,
- 0.4 g Na_2_MoO_4_.2H_2_O,
- 0.3 g CoCl_2_.6H_2_O
- 0.1 g CuSO_4_.5H_2_O
- 0.1 g KI
- 15 g EDTA

**The **vitamins stock solution** contained (L^-1^):

- 50 mg biotin
- 200 mg p-aminobenzoic acid
- 1 g nicotinic acid
- 1 g Ca-pantothenate
- 1 g pyridoxine-HCl
- 1 g thiamine-HCl
- 25 g myo-inositol

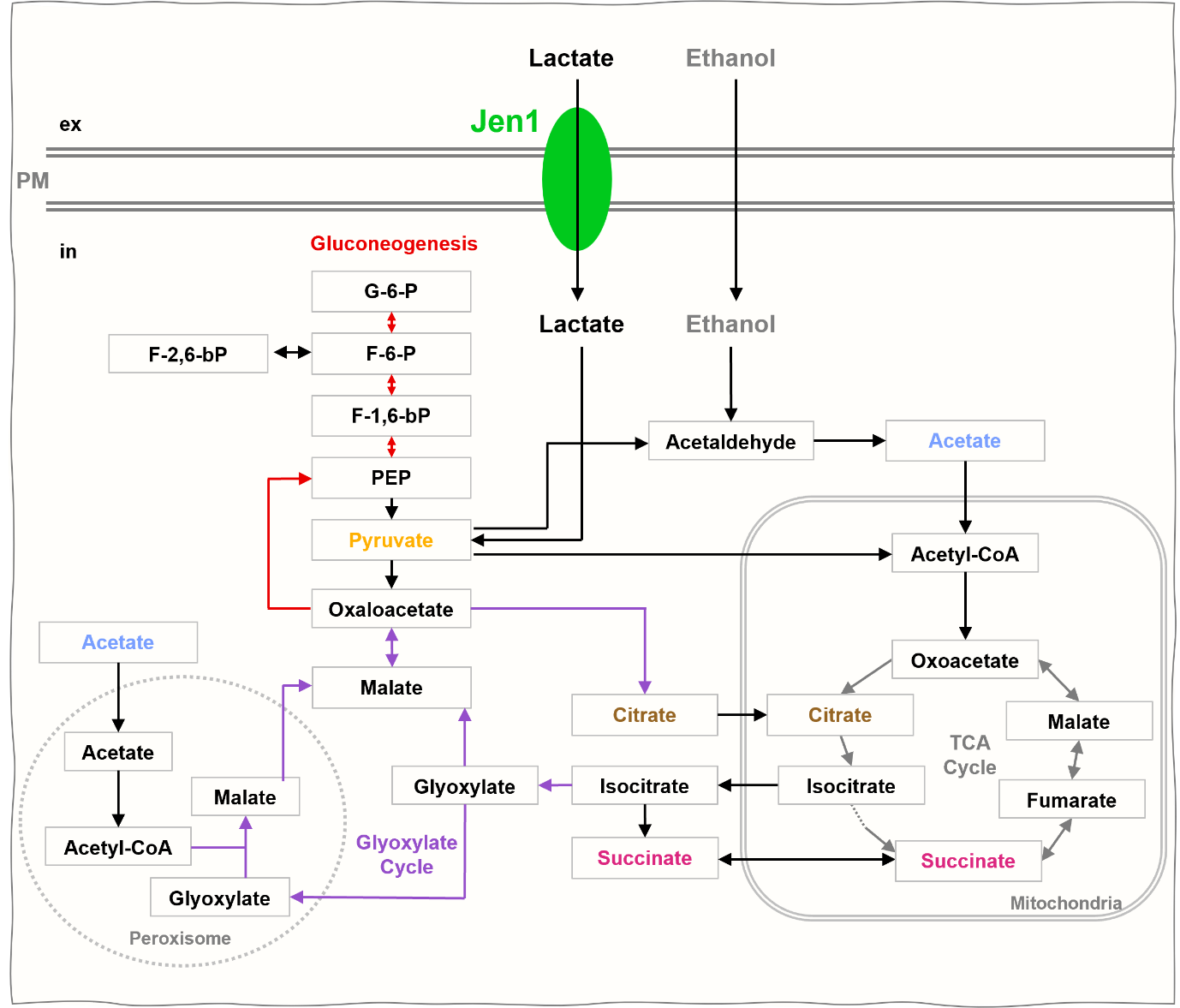

Fig. S4 Schematic representation of the main metabolic pathways involved in the utilization of lactate and ethanol as carbon sources in the yeast *S. cerevisiae*.

The TCA (tricarboxylic acid) cycle is represented in grey; the Glyoxylate cycle in purple and Gluconeogenesis in red. Some steps in the TCA cycle are omitted (adapted from [1]).
